# De novo deletions and duplications at recombination hotspots in mouse germlines

**DOI:** 10.1101/2020.06.23.168138

**Authors:** Agnieszka Lukaszewicz, Julian Lange, Scott Keeney, Maria Jasin

## Abstract

Numerous DNA double-strand breaks (DSBs) arise during meiosis to initiate homologous recombination. These DSBs are usually repaired faithfully, but here we uncover a new type of mutational event in which deletions form via joining of ends from two closely spaced DSBs (double cuts) within a single hotspot or at adjacent hotspots on the same or different chromatids. Deletions occur in normal meiosis but are much more frequent when DSB formation is dysregulated in the absence of the ATM kinase. Events between chromosome homologs point to multi-chromatid damage and aborted gap repair. Some deletions contain DNA from other hotspots, indicating that double cutting at distant sites creates substrates for insertional mutagenesis. End joining at double cuts can also yield tandem duplications or extrachromosomal circles. Our findings highlight the importance of DSB regulation and reveal a previously hidden potential for meiotic mutagenesis that is likely to affect human health and genome evolution.

## Introduction

Meiotic recombination is essential for faithful genome transmission in most sexually reproducing organisms (Hunter, 2015). Recombination is initiated by the programmed formation of DNA double-strand breaks (DSBs) by the SPO11 protein (Keeney, 2001). SPO11 preferentially cuts within narrow genomic regions called hotspots (Baudat et al., 2013; de Massy, 2013; Petes, 2001). Of all DSB hotspots (~15,000 defined in mouse), only a tiny fraction (~250-350) experiences a DSB in any single meiosis (Cole et al., 2012; Lange et al., 2016; Li et al., 2019; Tock and Henderson, 2018). DSBs are also nonrandomly dispersed along chromosomes to ensure homolog pairing and recombination (Keeney et al., 2014). This quantitative and spatial regulation of hotspot usage involves hierarchical and combinatorial levels of control (Lange et al., 2016; Pan et al., 2011; Yamada et al., 2017), with the ATM kinase playing a central role (Cooper et al., 2016; Lukaszewicz et al., 2018).

ATM, known for its role in the DNA damage response in mitotic cells (Shiloh, 2006), is activated by meiotic DSBs to limit DSB numbers and to control hotspot usage genome-wide (Lange et al., 2011; Lange et al., 2016). Control of DSB numbers by ATM and its orthologs is conserved, although ATM-deficient mice have the largest reported increase in DSB levels (e.g., ~10 fold compared with ~2 fold in budding yeast deficient for the ATM ortholog Tel1) (Checchi et al., 2014; Kurzbauer et al., 2021; Lukaszewicz et al., 2018; Tian and Loidl, 2018). Control of genome-wide DSB distributions is also conserved (Mohibullah and Keeney, 2017). In yeasts, Tel1 controls DSB formation locally by suppressing double cutting, that is, simultaneous DSB formation at adjacent hotspots (Cooper et al., 2016; Fowler et al., 2018; Garcia et al., 2015). This suppression is strongest at closely-spaced hotspots (<10 kb), such that in the absence of Tel1, double cutting at hotspots separated by ~2 kb is more frequent than expected by chance and ~10-fold higher than in wild type. The loss of this Tel1-mediated DSB suppression is thought to contribute to the global increase in DSBs (Garcia et al., 2015; Mohibullah and Keeney, 2017). In mouse, double cutting at adjacent hotspots has not been demonstrated, but potentially may be even more likely in the absence of ATM, considering the extremely elevated DSB numbers. Accordingly, we previously suggested that ATM may inhibit multiple nearby DSBs (Lange et al., 2011; Lange et al., 2016).

Normally, most meiotic DSBs are repaired by recombination between homologous chromosomes. While nonhomologous end-joining (NHEJ) is a robust mitotic DSB repair pathway in mice (Chang et al., 2017), it is thought to be suppressed in meiosis (Kim et al., 2016). However, NHEJ can occur in mouse spermatocytes when DSBs are induced by irradiation (Ahmed et al., 2010; Ahmed et al., 2018; Enguita-Marruedo et al., 2019), and in *C. elegans* in mutants in which DSB processing (Girard et al., 2018; Lemmens et al., 2013; Yin and Smolikove, 2013) or recombination (Macaisne et al., 2018; Martin et al., 2005) is compromised.

The consequences of deregulated meiotic DSB formation are not fully understood. We hypothesized that, in the absence of ATM, formation of DSBs that are too numerous or too closely spaced interferes with the meiotic recombination program. This in turn might elicit the use of other repair pathways, thus compromising genome integrity. We reasoned that mutagenic NHEJ in particular might become more prevalent because of resection defects known to occur in the absence of ATM (Cannavo et al., 2018; Joshi et al., 2015; Mimitou et al., 2017; Paiano et al., 2020; Yamada et al., 2020). Moreover, the loss of ATM may cause DSBs to form at similar locations on multiple chromatids — due to the increase in DSB numbers and/or the loss of spatial control — and thus promote NHEJ by damaging the templates needed for recombinational repair.

We tested this hypothesis by asking whether multiple DSBs form at adjacent hotspots in the absence of ATM in mouse spermatocytes and whether they are repaired aberrantly to delete the intervening sequences. Further, we asked whether multiple DSBs leading to deletions can occur within an individual hotspot. In the course of these experiments, we uncovered novel DSB (mis)repair products that provide insight into the consequences of loss of DSB control during meiosis.

## Results and Discussion

### Hidden potential of the meiotic genome to form deletions

We first catalogued genomic locations at risk for deletion by identifying pairs of closely-spaced hotspots in nucleotide-resolution DSB maps obtained by sequencing SPO11-bound oligonucleotides (oligos), a by-product of DSB formation (Lange et al., 2016). We found that numerous DSB hotspots are separated by a short genomic distance (**Fig. 1A**). For example, ~6% of 13,436 autosomal SPO11-oligo hotspots in wild-type mice are located within 5 kb of another hotspot, and this percentage nearly doubles (~10%) in *Atm*^−/−^ mice, which have more hotspots called (20,417) (**Fig. S1A**). Most hotspots in *Atm*^−/−^ mice are shared with those in wild type, although a significant fraction are newly called because the few SPO11 oligos that cluster at these sites in wild type are greatly increased in the mutant (Lange et al., 2016) (**Fig. S1A**). Newly called hotspots often emerge near shared hotspots, resulting in an increased number of closely-spaced hotspots in *Atm*^−/−^ (**Fig. S1B**). Moreover, in addition to the number, the strength of the shared hotspot pairs also disproportionately increases in *Atm*^−/−^ (**Fig. S1C**), at least in part due to the particularly large increase in DSBs at weaker hotspots (Lange et al., 2016). Thus, while adjacent DSBs may normally be suppressed, this hidden potential for DSBs at adjacent hotspots may be revealed in *Atm*^−/−^.

**Figure 1.**
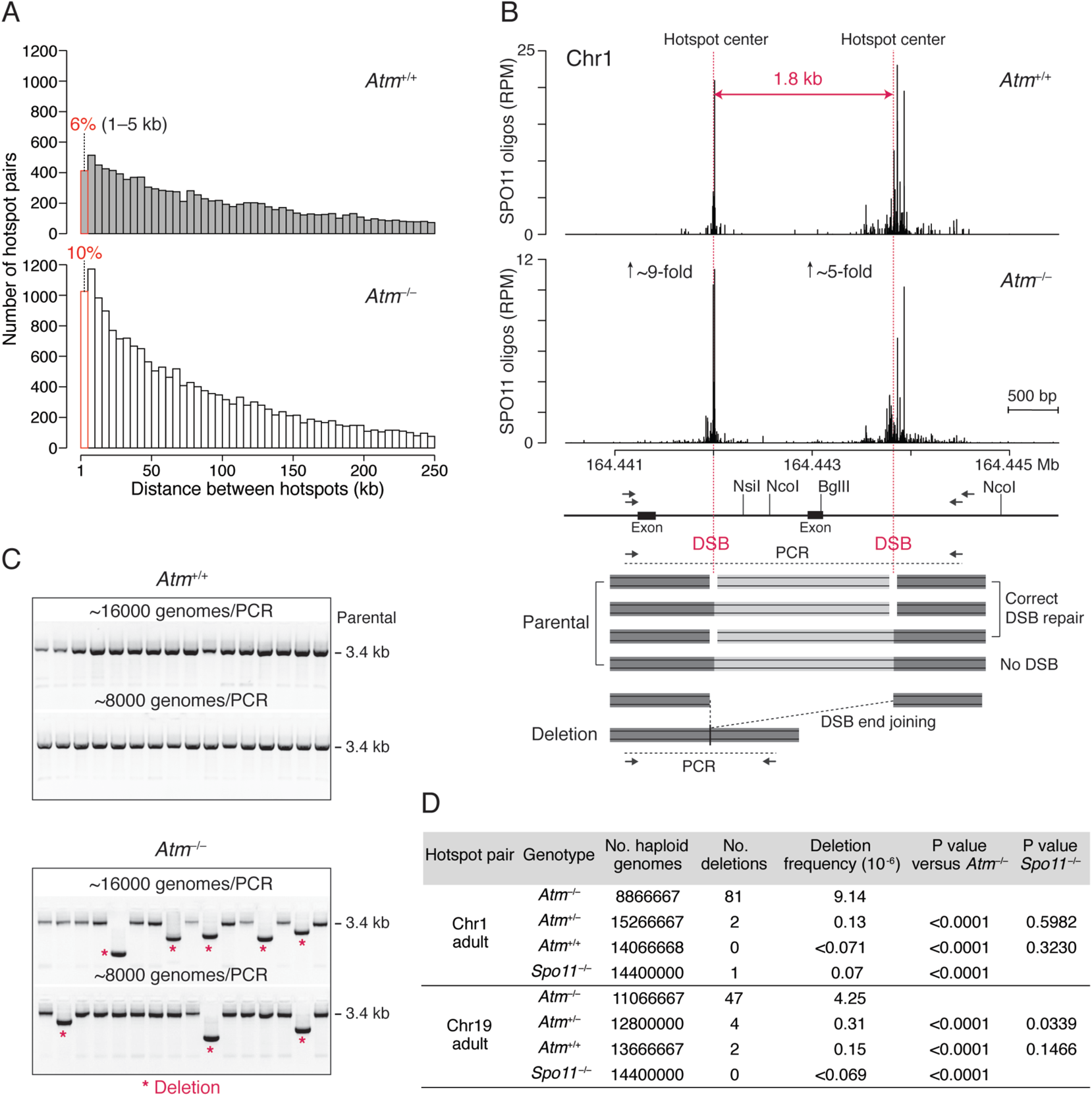
Meiotic deletions at adjacent DSB hotspots. **(A)** Distribution of distances separating adjacent SPO11-oligo hotspots on autosomes. The hotspot calling algorithm (Lange et al., 2016) merges hotspots separated by edge-to-edge distances of <1 kb. Hotspot pairs separated by >250 kb are not shown. See also **Fig. S1**. **(B)** Strategy to detect deletions between adjacent hotspots. Top panel, SPO11-oligo maps for the hotspot pair on Chr1 (RPM, reads per million). Genomic coordinates (MM10 assembly) are shown below the graphs. The estimated fold increase in absolute SPO11-oligo frequency (RPM × 11.3) is indicated for each hotspot. Vertical pink lines indicate hotspot centers. Lower panel, scheme of the PCR assay. Nested PCR across the DSB hotspot pair is carried out with the indicated primer pairs (arrows). Prior to amplification, genomic DNA is optionally digested with restriction enzymes to reduce amplification of parental products. The *Atp1b1* exon located between the two hotspots is removed in deletions. **(C)** Detection of deletions by agarose gel electrophoresis. Representative gels for the Chr1 hotspot pair are shown. Each lane is a PCR seeded with the indicated number of haploid genomes. **(D)** Deletion frequencies. Three to seven mice of each genotype per hotspot pair were analyzed, listed in **Table S1**. P values are from the Chi-square test.

To test for deletions, we selected DSB hotspot pairs that are shared between wild type and mutant and that contain strong hotspots to increase the chance of simultaneous DSB formation. The chosen hotspot pairs on Chr1 and Chr19 are separated by <2 kb, with hotspots that individually rank in the top quartile of all hotspots in both wild type and *Atm*^−/−^ (**Fig. 1B, S1D, S2A**). We tested for deletions between the two hotspots in each pair by performing nested PCR on testis DNA. For both pairs, concurrent DSB formation at the two hotspot centers with precise joining of the DNA ends would give rise to 1.8-kb deletions (**Fig. 1B, S2A**); in practice, deletions would vary in size depending on the location of SPO11 cleavage in any particular cell. To suppress amplification of parental DNA, genomic DNA was digested with restriction enzymes prior to amplification in some experiments. Nested PCR was performed with input testis DNA equivalent to ~8,000–16,000 haploid genomes per well for a total of ~10–15 million haploid genomes per genotype (**Table S1**).

Smaller PCR products indicative of deletions between the two DSB hotspots were observed in *Atm*^−/−^ (**Fig. 1C, S2B**). In adult *Atm*^−/−^ mice, the frequencies of deletions at the Chr1 and Chr19 hotspot pairs were 9.14 × 10^-6^ and 4.25 × 10^-6^, respectively, per haploid genome (**Fig. 1D**; **Table S1**). Deletions were also detected in ATM-proficient cells but were rare. At the Chr19 hotspot pair, the frequencies were 0.15 × 10^-6^ and 0.31 × 10^-6^ in wild type and *Atm*^+/−^, respectively, reflecting a total of just six events, while no deletions were observed in *Spo11*^−/−^. For the Chr1 hotspot pair, only two deletions were observed in ATM-proficient cells, although one was also observed in *Spo11*^−/−^.

Similar results were obtained in juvenile testes at both hotspot pairs at 15 days post partum, when wild-type and mutant testes have a similar distribution of prophase I stages. This finding indicates that the increase in deletion frequency is not due to the different cellular composition of mutant testes caused by prophase I arrest of *Atm*^−/−^ spermatocytes (Barlow et al., 1998; Xu et al., 1996) (**Table S1**).

Furthermore, we tested for the presence of deletions in *Atm*^+/−^ and *Atm*^+/+^sperm DNA. We captured 7 deletions in *Atm*^+/−^ at the Chr19 hotspot pair, for a frequency 1.09 × 10^-6^ per haploid genome, although none from *Atm*^+/+^ (≤0.31 × 10^-6^; **Table S1**). This demonstrates that meiotic deletions can be transmitted into sperm.

It should be noted that testis DNA is derived from germ cells at different stages of development as well as from some somatic cells, such that the germline deletion frequencies we report for testes are underestimated for all genotypes. Even so, meiotic recombination at a well-studied hotspot that has a similar strength as the individual hotspots examined here occurs at a substantially higher frequency (two to three orders of magnitude: 5.8 × 10^-3^ noncrossovers per sperm genome (Cole et al., 2010)) than the deletion frequency we observed at the hotspot pairs in the absence of ATM. However, considering the large number of hotspots located close to one another, the potential to form deletions in meiosis is substantial.

### Double cutting in the absence of ATM

To map deletion breakpoints, we sequenced deletion products from *Atm*^−/−^ adults and juveniles: 134 from the Chr1 hotspot pair and 58 from the Chr19 hotspot pair (**Fig. 2A, S2C**; **Table S2**). The distribution of deletion breakpoints resembled the distribution of SPO11 oligos at both hotspots, with breakpoints clustered around hotspot centers (**Fig. 2B,C, S2D,E**). Smoothed breakpoint and SPO11-oligo profiles displayed similar shapes, except that breakpoint profiles were slightly wider with more prominent smaller peaks (**Fig. 2D, S2F**). We defined the breakpoint center within each hotspot as the position of the smoothed peak in the breakpoint density, in the same manner as hotspot centers were defined (Lange et al., 2016). Breakpoint centers closely matched all four hotspot centers (within 13 to 39 bp). These findings strongly indicate that deletion formation is preceded by concurrent SPO11 cleavage at the two hotspots, providing the first evidence for double cutting at adjacent DSB hotspots in mouse.

**Figure 2.**
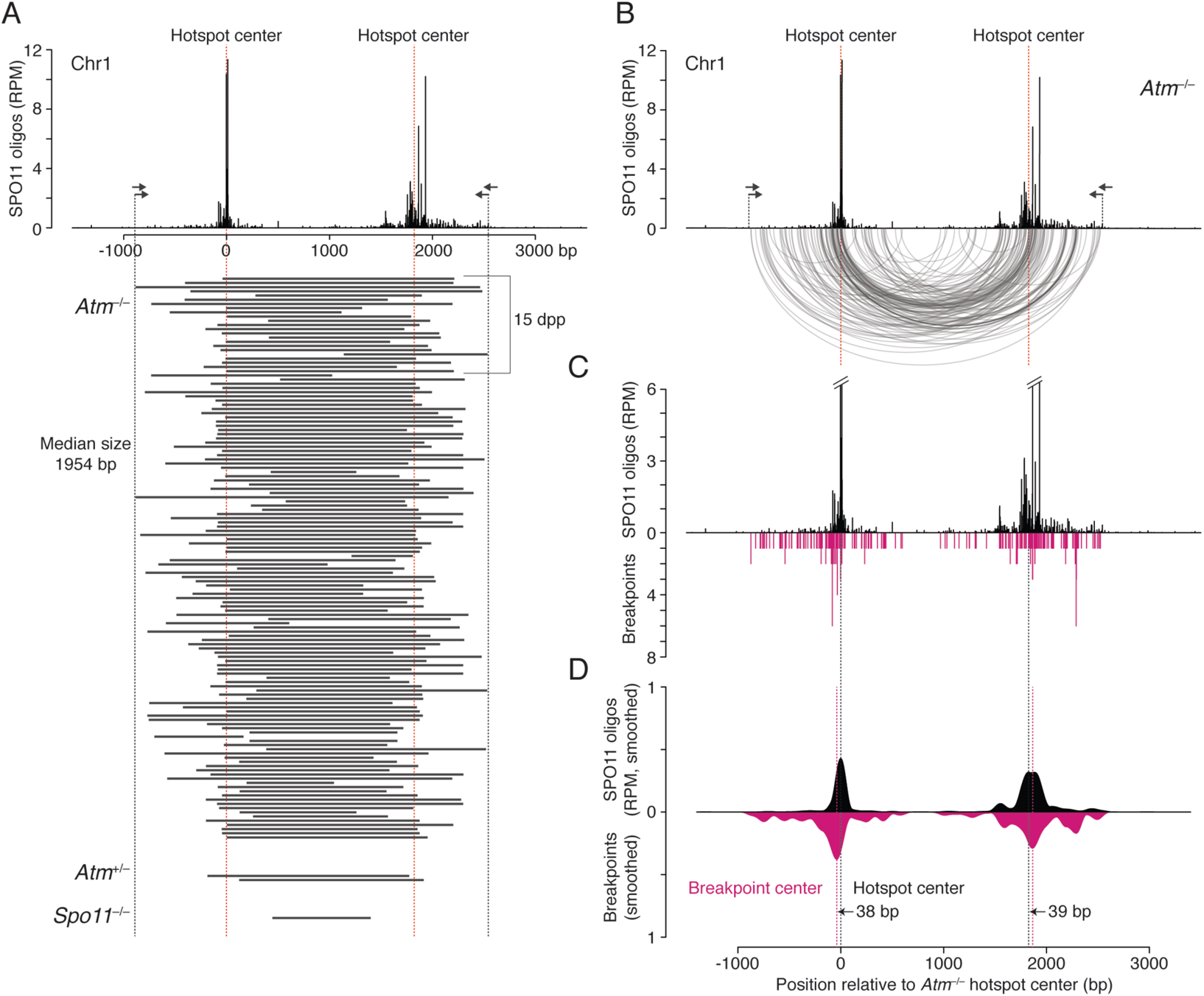
Deletion breakpoints at the Chr1 hotspot pair. **(A)** Deleted DNA sequences at the Chr1 hotspot pair. Deletions (black bars) were combined from adult and juvenile testes (dpp, days post partum). **(B)** Arc diagram of deletions in *Atm*^−/−^ at the Chr1 hotspot pair. Arcs link the breakpoints of each deletion (134 deletions total). Both deletion breakpoints often map close to the hotspot centers. See also **Fig. S3**. **(C)** Individual breakpoints cluster around the hotspot centers but are also observed to spread. **(D)** Smoothed breakpoint and SPO11-oligo profiles display similar shapes (201-bp Hann filter for smoothing). Offsets between centers of hotspots and breakpoint distributions are indicated.

The median deletion lengths were similar to the distances between the DSB hotspot centers (Chr1: 1954 bp vs 1825 bp; Chr19: 1892 bp vs 1763 bp), as expected from SPO11 cleavage of both hotspots and joining of unprocessed or minimally processed DNA ends. Accordingly, many deletions had breakpoints within the narrow (251 bp) central zones around the hotspot centers that contained the majority (>60%) of SPO11 oligos: Nearly half of the deletions had one breakpoint within the central zone of one hotspot (Chr1, 41%; Chr19, 45%) and about a fifth had both breakpoints within the central zone (Chr1, 21%; Chr19, 19%) (**Fig. S3A**). While the number of deletions with both breakpoints within the zone was lower than predicted from the SPO11-oligo counts (Chr1, 65%; Chr19, 48%; **Fig. S3B**), it was notable that breakpoints outside the zone were not evenly distributed. Instead, some clusters could be observed that overlapped with, but were more prominent than, SPO11-oligo clusters (**Fig. 2B–D, S2D–F**). The overlap of breakpoints and SPO11 oligos implies that DSB ends are usually joined close to the initial SPO11 cleavage sites, whereas the wider distribution of deletion breakpoints may reflect preferential deletion formation for DSBs further from the hotspot center, DNA end processing prior to joining, or even multiple DSBs within the same hotspot (see below).

In principle, deletions that we observe could arise from single DSBs at one hotspot that are asymmetrically resected. However, this seems highly unlikely, given that both breakpoints usually map in the vicinity of separate hotspots, even with the omission of the restriction digest step (**Fig. S4**). Moreover, end-joining at single, unprocessed DSBs is predicted to be error-free due to the presence of 2-bp 5′ overhangs after SPO11 cleavage (Hatkevich et al., 2021; Yamada et al., 2020). Although our assay is not designed to detect small deletions at a single hotspot, we address this question below.

Notably, the frequency of deletions in *Atm*^−/−^ at both hotspot pairs is ~100 times lower than the frequency of double cutting estimated from SPO11-oligo read counts with the assumption of independent cutting (**Table S3**). While our assay may not detect all deletions, it seems likely that only a subset of double cuts undergoes end joining, while the others are correctly repaired, misrepaired by other mechanisms, or not repaired at all, given that *Atm*^−/−^ spermatocytes accumulate unrepaired DSBs (Barchi et al., 2005).

In wild type and *Atm*^+/−^, several among the small number of deletions recovered had breakpoints that were identical to those observed in *Atm*^−/−^, while the one deletion obtained from *Spo11*^−/−^ had both breakpoints comparatively far from the hotspot centers (**Fig. S5**). However, most of the deletion breakpoints in wild type and *Atm*^+/−^ were not within the narrow central zone (251 bp) of SPO11 oligos around the hotspot centers. For example, 12 deletions (86%) at the Chr19 hotspot pair had both breakpoints outside this zone. Although more spread out, breakpoints still appeared to often cluster where SPO11 oligos clustered (**Fig. S5**). These observations suggest that in the presence of ATM, deletions may be more prone to occur for DSBs formed outside of hotspot centers or the DSBs involved in the deletion may be more processed.

### Double cutting can involve one or multiple chromatids leading to distinct mutational outcomes

The meiotic deletions we observe could arise from two DSBs on a single chromatid or DSBs on two chromatids. Moreover, double cutting produces four DNA ends, with the potential to give rise to other mutational outcomes besides deletions. In particular, DSBs on two chromatids can generate tandem duplications as well as deletions, while two DSBs on one chromatid can release a DNA fragment with the potential to circularize (**Fig. 3A**). Finally, DSBs on homologous chromosomes may provoke inter-homolog end-joining. We investigated each of these possibilities.

**Figure 3.**
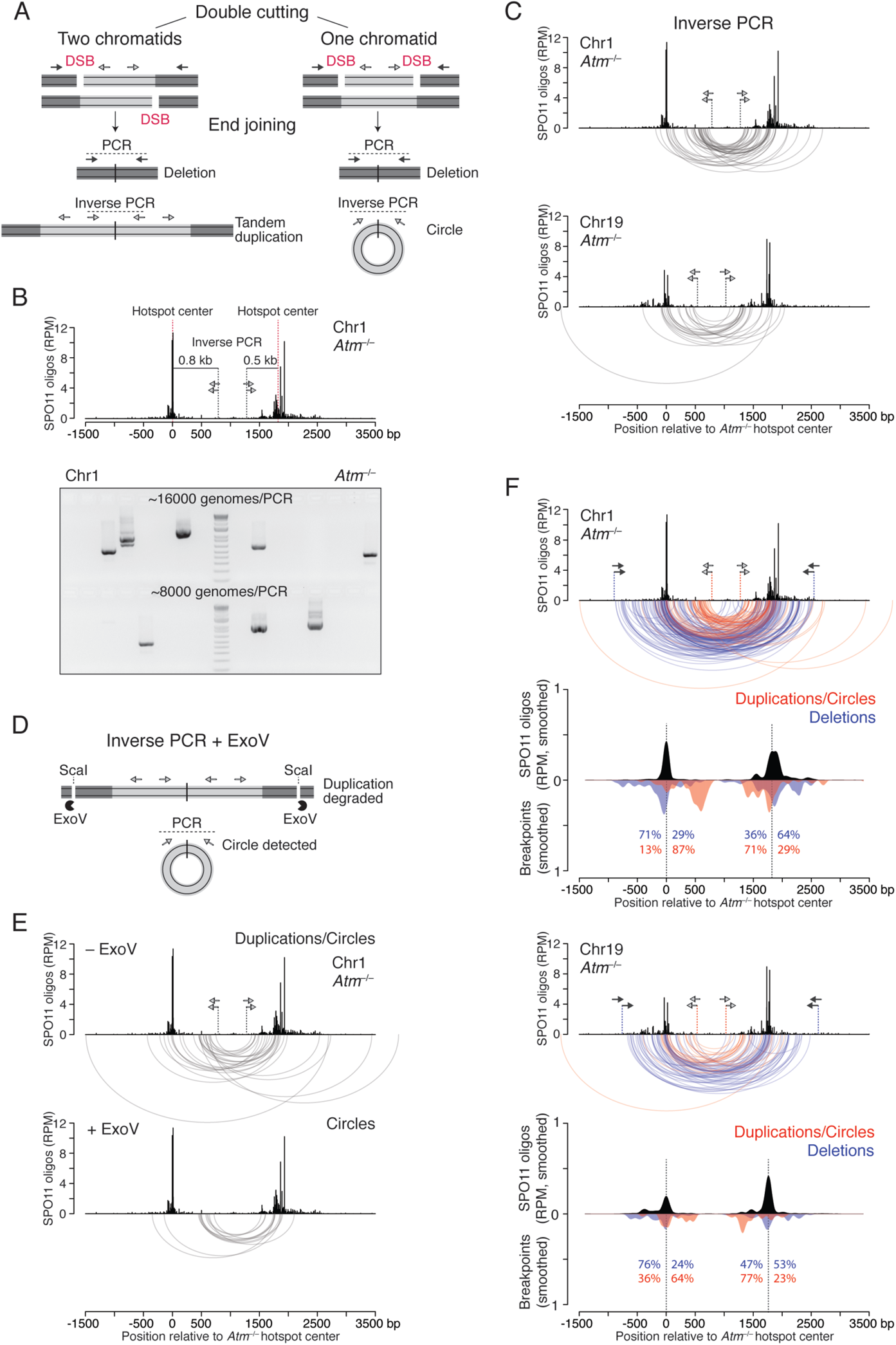
Tandem duplications and circular DNA from double cutting. **(A)** Predicted end-joining products from double cutting in one or two chromatids. Lighter grey indicates DNA segments between two DSBs at adjacent hotspots. PCR with primers outside of the two DSBs (dark grey arrows) detects deletions. Inverse PCR, with primers inside of the two DSBs (light grey arrows), detects both tandem duplications and circles, which generate identical PCR products. **(B)** Inverse PCR assay at the Chr1 hotspot pair. Top panel, nested PCR is carried out with the indicated primer pairs (light gray arrows). A 1.3-kb PCR product would arise from a tandem duplication or circular DNA from end joining of double cuts precisely at the two hotspot centers. Lower panel, inverse PCR products detected in *Atm*^−/−^ testis DNA. PCR reactions were seeded with the indicated number of haploid genomes. **(C)** Arc diagrams of breakpoints for duplications/circles at the Chr1 (42 events) and Chr19 (22 events) hotspot pairs in *Atm*^−/−^. **(D)** Strategy to detect only circular DNA products by inverse PCR. ExoV is used to degrade linear DNA containing tandem duplications. ScaI produces entry sites for ExoV degradation close to tandem duplications. See also **Fig. S6B**. **(E)** Arc diagrams of duplications/circles (27 events) and circles (13 events) at the Chr1 hotspot pair, from *Atm*^−/−^ testis DNA treated without and with ExoV, respectively. Equal numbers of haploid genomes were tested in both cases (**Fig. S6C**). **(F)** Distinct breakpoint profiles for duplications/circles and deletions at the Chr1 and Chr19 hotspot pairs in *Atm*^−/−^. Arc diagrams are shown for each hotspot pair, with duplications/circles (red arcs; 69 events) and deletions (blue arcs) overlaid. Note that duplication/circle arcs are often smaller than deletion arcs and that duplication/circle breakpoints occur more often in between the two hotspots. Below the arc diagrams are smoothed breakpoint and SPO11-oligo profiles (201-bp Hann filter). Duplication/circle profiles are normalized to the number of deletions to better compare the two profiles. Percentages indicate the fraction of breakpoints mapping to the left or right of the hotspot centers. See also **Fig. S6D**.

#### Tandem duplications and circular DNA

We used outward-directed (inverse) PCR, which would give a product from either a tandem duplication or circular DNA (duplication/circle) arising from double cutting (**Fig. 3A**). In *Atm*^−/−^ testis DNA, duplication/circle products were readily detected (**Fig. 3B**) at a frequency of 2.17 × 10^-5^ per haploid genome for the Chr1 hotspot pair (42 events) and 2.20 × 10^-5^ for the Chr19 pair (22 events) (**Fig. S6A**). (We excluded a fraction of products that we were unable to fully sequence due to the presence of palindromes at the junctions, the origin of which is unclear; see **Fig. S6A**.) This frequency is higher than what we obtained for deletions. Because duplications/circles and deletions are predicted to be reciprocal products, these differences suggest that deletions are underdetected. Alternatively, joining of ends participating in duplication/circle formation is somewhat more efficient than that of deletions. Both hotspot pairs showed similar breakpoint distributions, with some breakpoints clustering at the most prominent SPO11-oligo peaks while others clustered between the two hotspots (**Fig. 3C**) (see below).

Next, we aimed to distinguish circular DNA junctions from those of duplications to determine if both events occur. To this end, genomic DNA was incubated with *E. coli* exonuclease V (also known as RecBCD) prior to inverse PCR to specifically degrade linear DNA containing duplications while leaving circles intact (**Fig. 3D, S6B**). We cleaved DNA with a restriction enzyme with sites flanking the hotspot pair to further ensure ExoV degradation, which was confirmed with a control PCR (**Fig. S6B**). Using this approach at the Chr1 hotspot pair, we recovered 13 products from reactions with ExoV, which are predicted to arise only from circles, and a significantly larger number (27) without ExoV (P = 0.0269, Chi-square test; **Fig. 3E, S6C**). This result provides evidence that tandem duplications and circles both occur in *Atm*^−/−^ testis DNA, and in roughly similar numbers. A few presumptive tandem duplications were also detected in *Atm^+/+^* sperm DNA (**Fig. S6C**), although further investigation is required.

The circle and duplication/circle breakpoint distributions were similar, indicating that end joining positions are not dramatically different within the hotspot pair despite involving DSBs on different numbers of chromatids (**Fig. 3C, 3E**). Distances between the breakpoints, which indicate the duplication length or the circle size, were likewise similar (Chr1: circles, 1267 bp median; duplications/circles, 1287 bp median). In contrast, these distances were significantly smaller than those of deletions (1954 bp median; P < 0.0001, unpaired Mann-Whitney test). Similar findings were obtained for the Chr19 hotspot pair (median deletion size was 1892 bp, compared with 1348 bp for duplications/circles; P < 0.0001, Mann-Whitney test; **Fig. 3C**). This difference was because duplication/circle breakpoints mapped between hotspot centers (64–87%), whereas the majority of deletion breakpoints mapped outside (53–76%, **Fig. 3F**). These shifts were highly significant (**Fig. S6D**) and are consistent with multiple DSBs within hotspots and/or joining of partially resected DSB ends, because both scenarios would yield larger deletions and smaller duplications and circles (**Fig. S6E**).

Interestingly, for duplications/circles only one of the two major breakpoint peaks in each pair was close to the hotspot center. The other was shifted ~500 bp inward (Chr1 left hotspot, Chr19 right hotspot, **Fig. 3F**). This shift may be the result of frequent double cuts within these two hotspots, with one cut at the hotspot center and the other ~500 bp away. Supporting this, a small cluster of SPO11-oligos is observed at this position in the Chr19 right hotspot (**Fig. 3F**) and is also seen in the Chr1 left hotspot when SPO11-oligos mapping to repeats are included (**Fig. S6F**).

Overall, our results provide evidence for multiple mutational outcomes arising from double cutting and end joining in the absence of ATM. Moreover, as circles and tandem duplications are not predicted to arise from a single DSB, their occurrence provides strong evidence for double cutting at two DSB hotspots on either one or two chromatids.

#### Joining between homologous chromosomes

To investigate whether deletions involve one or both homologs, we tested the Chr1 hotspot pair in B6 × DBA/2J (DBA) F1 hybrid males, which have a high polymorphism density across this region (**Fig. 4A**). Primers were shifted to amplify a larger fragment to aid in identifying the parental chromosome(s) involved in the deletion.

**Figure 4.**
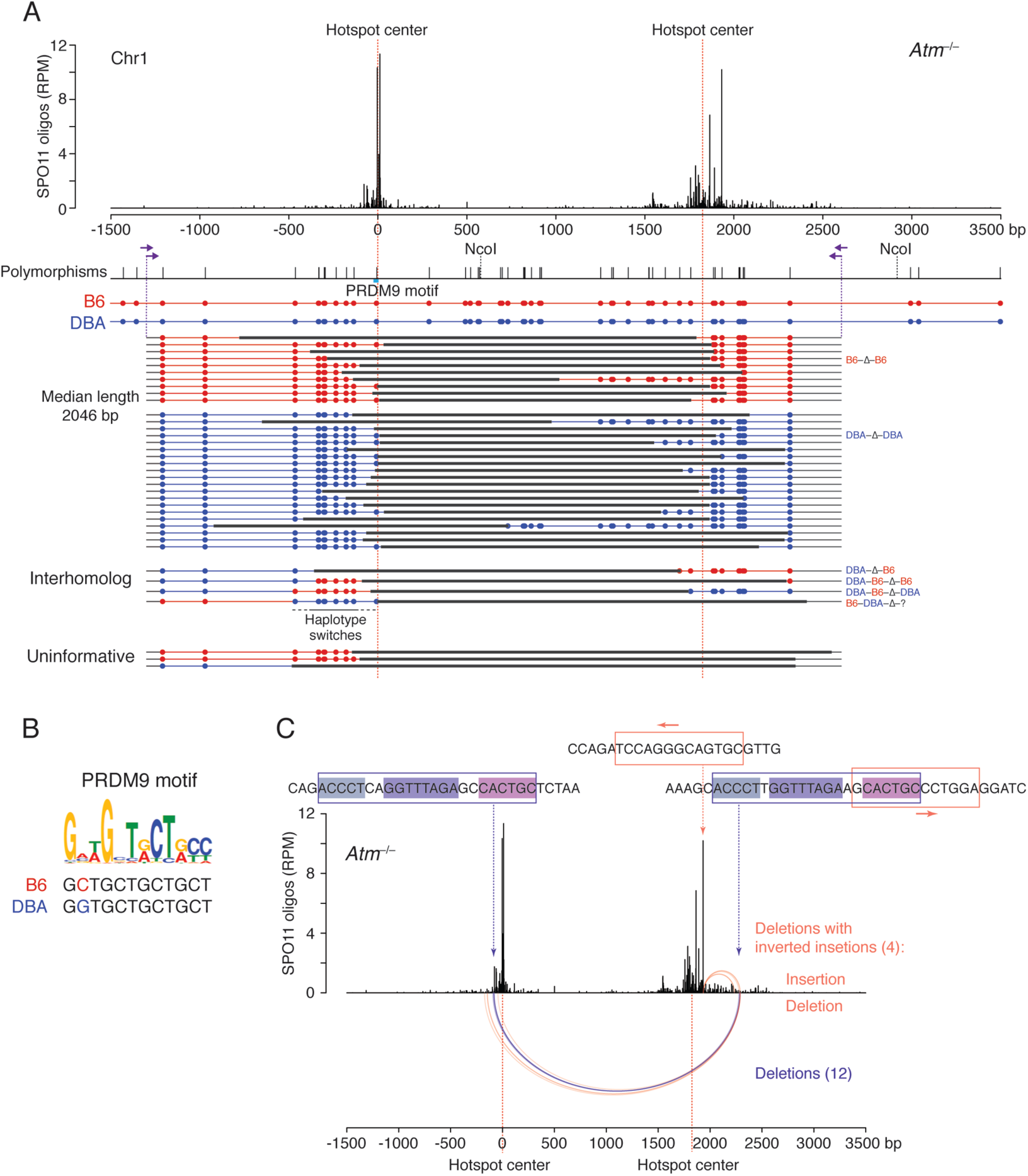
Meiotic deletions within and between chromosomes. **(A)** Deletions at the Chr1 hotspot pair in *Atm*^−/−^ B6 × DBA F1 hybrid males. Nested PCR was performed with the indicated primers (arrows) to identify polymorphisms within the deletion products derived from B6 (red) or DBA (blue). Three deletions were uninformative because the involved chromosome(s) is uncertain. **(B)** Predicted PRDM9 recognition sequences at the left hotspot in B6 and DBA, aligned to the PRDM9 motif from SPO11-oligo analysis (Lange et al., 2016). **(C)** Recurrent deletion within the Chr1 hotspot pair at an imperfect repeat. Twelve simple deletions in *Atm*^−/−^ map to this repeat at one of the three internal identical repeats of 5, 8, and 6 bp (violet arcs). Four other deletions with inverted insertions also involve this imperfect repeat as well as a 13-bp inverted microhomology at the hotspot center, with the insertions indicated above the deletions (orange arcs). See also **Fig. S8A, S10B**.

We recovered 37 deletions from *Atm*^−/−^ testis DNA for a frequency of 4.20 × 10^-6^ per haploid genome, but none in the presence of ATM (<0.16 × 10^-6^) (**Table S1**). The distributions of deletion sizes (median 2046 bp) and breakpoints were broadly similar to those from B6 inbred mice (**Fig. 4A, S7A**). When we analyzed polymorphisms, 30 of the 34 informative deletions (88%) appeared to have originated within one chromosome: 10 on B6 and 20 on DBA (**Fig. 4A**). The higher frequency of deletions on the DBA chromosome might be associated with a polymorphism in the PRDM9 binding site at the left hotspot (**Fig. 4A,B**), which could positively affect DSB formation. Thus, the preponderance of deletions arise within one chromosome, involving either one chromatid or both sister chromatids.

Strikingly, however, we also captured four deletions that contained sequences from both homologs (**Fig. 4A**). In one, the left hotspot from DBA was joined to the right hotspot from B6 (DBA-Δ-B6). In the other three more complex events, a haplotype switch occurred adjacent to one of the deletion breakpoints (DBA-B6-Δ-B6; DBA-B6-Δ-DBA, and B6-DBA-Δ-?, where the question mark indicates that the genotype was uninformative at the right end). In these haplotype switches, 6–8 polymorphisms were incorporated, encompassing up to 500 bp (**Fig. 4A**).

The recovery of deletions involving interhomolog interactions shows that DSBs can occur at nearby locations on both homologs in the same cell, at least in the absence of ATM. Moreover, the DSB ends generated on the two homologs are capable of joining. Thus, although homolog synapsis is greatly impaired in the absence of ATM (Barlow et al., 1998; Xu et al., 1996), homolog interactions still occur. Cutting of both homologs at the same hotspot has been inferred from patterns of strand exchange in yeast tetrads (Anderson et al., 2015; Marsolier-Kergoat et al., 2018; Zhang et al., 2011).

Further, the three complex interhomolog events suggest that three or more DSBs had occurred on the two homologs. One possible mechanism for the haplotype switch at a deletion breakpoint is offset DSB formation on both homologs within the same hotspot, with attempted gap repair that is aborted by the DSB in the template, followed by joining to a DSB at the other hotspot (**Fig. S7B**). Alternatively, double cutting within one hotspot on one homolog might release a DNA fragment that then inserts at the DSB in the other homolog (**Fig. S7C**). Irrespective of the precise mechanism, however, these results indicate that loss of ATM can cause damage to multiple chromatids, leading to deleterious types of repair.

### Sites of recurrent deletion breakpoints in the absence of ATM

Most deletions between hotspots from *Atm*^−/−^ mice were unique: 87% (161/185) of events on Chr1 were observed only once, as were 97% (56/58) for Chr19 (**Table S2**). Microhomologies were present at the breakpoints in 75% of all deletions but were generally short, with a median of 1 bp for both the Chr1 and Chr19 deletions (**Fig. S8A**). Thus, substantial microhomology is not required for joining. However, the 24 recurring deletions on Chr1 had longer microhomologies with a median 6 bp (**Fig. S8B**). Twelve of the recurring deletions arose within a 22/23-bp imperfect sequence repeat shared between the two hotspots, and eight of these involved an 8-bp microhomology within the repeat (**Fig. 4C, S8B**). Thus, the availability and distribution of short sequence repeats likely influences deletion outcomes.

The presence or absence of microhomologies within end-joining products may be indicative of the end joining pathway used and/or processing at DNA ends. For example, SPO11-bound DSB ends normally require endonucleolytic processing on one strand by MRE11 to release SPO11 oligos (Keeney et al., 1997; Lange et al., 2011; Neale et al., 2005). This may disfavor canonical nonhomologous end-joining and promote alternative pathways that utilize microhomology (Sfeir and Symington, 2015). However, as some DSBs remain unresected in the absence of ATM/Tel1 (Cannavo et al., 2018; Joshi et al., 2015; Mimitou et al., 2017; Paiano et al., 2020; Yamada et al., 2020), deletions may also arise from joining at or very close to SPO11 cleavage positions, which would involve alternative SPO11 removal mechanisms, such as TDP2 (Alvarez-Quilon et al., 2014; Johnson et al., 2019a; Paiano et al., 2020). Our findings are thus consistent with multiple end-joining mechanisms being employed to repair adjacent DSBs in the absence of ATM.

### Insertion of hotspot-derived fragments

Unexpectedly, we observed insertions in 11% of deletions from *Atm*^−/−^ mice (27/243; two deletions with two inserts, for 29 total inserts; **Table S2**). All but one of the inserted DNA fragments mapped to chromosome positions showing SPO11 activity, either the assayed hotspot pairs themselves or other hotspots. The insert that mapped to a sequence with no SPO11 oligos contained a CT-rich microsatellite repeat and shared significant microhomology with both deletion breakpoints (12 and 9 bp) (**Table S3**), suggesting a possible DNA synthesis-based insertion mechanism.

#### Ectopic insertions from other hotspots

Fifteen inserts of 15–349 bp mapped to other chromosome positions at hotspots or SPO11-oligo clusters that were below the hotspot-calling threshold (**Fig. 5A-C, S9, S10A**; **Table S4**). Microhomologies at both sides of inserts were often short (0 to 2 bp), similar to deletions without insertions (**Fig. 5D**; **Table S4**).

**Figure 5.**
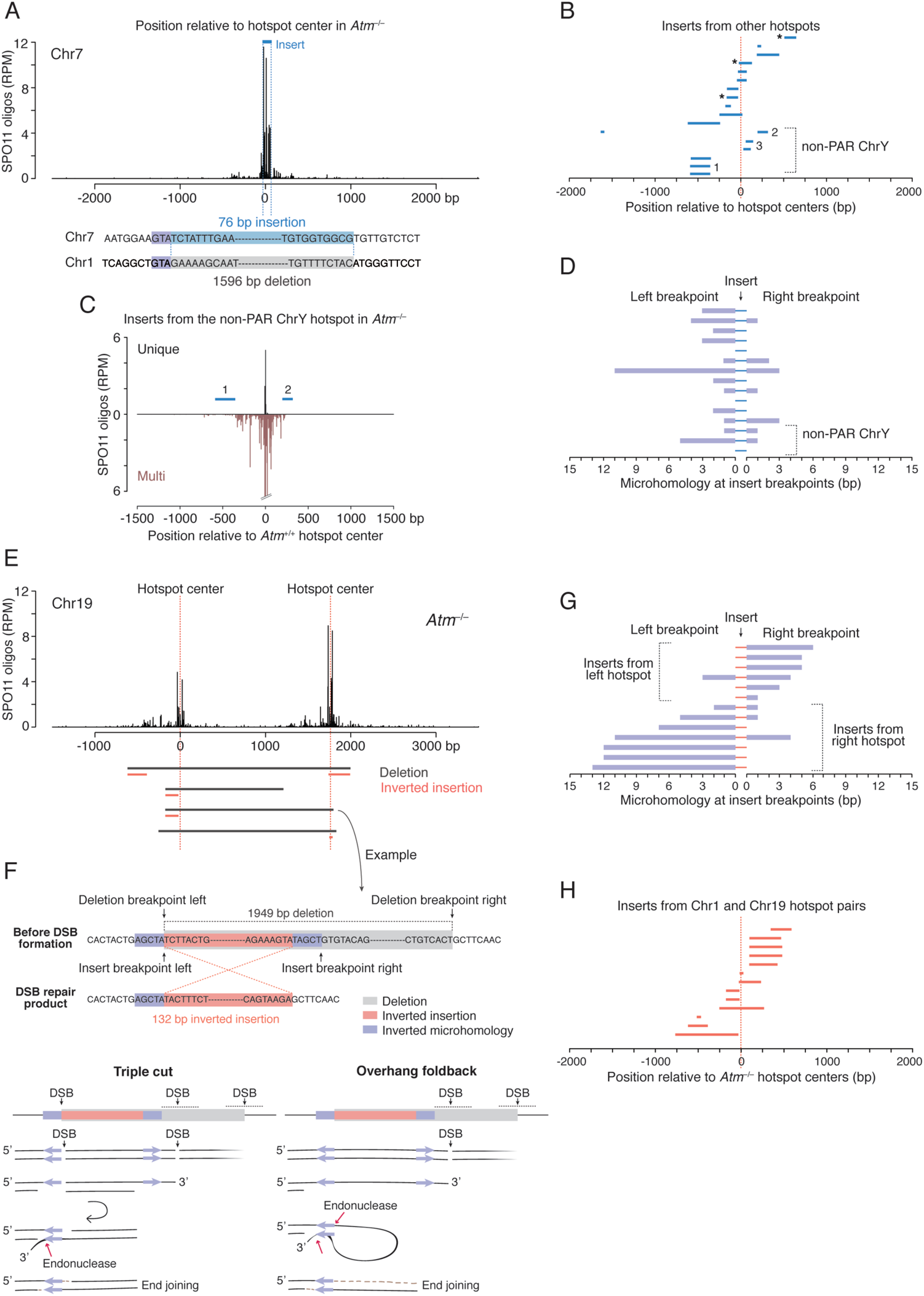
Insertions derived from DSB hotspots. **(A–D)** Ectopic insertions from other hotspots. **A**, Example of a deletion on Chr1 from *Atm*^−/−^ containing a 79-bp insertion derived from a hotspot on Chr7. **B**, Inserts from other hotspots map close to their respective hotspot centers. Three inserts map to SPO11-oligo clusters rather than called hotspots (asterisks). Inserts 1 and 3 from the non-PAR ChrY map to multiple hotspots. See also **Fig. S9, S11** and **Table S4**. **C**, Non-PAR ChrY hotspot to which inserts 1 and 2 can be mapped. Some SPO11 oligos can be uniquely mapped to this hotspot, but, as it is one copy of a family of repeats, numerous other SPO11 oligos can be mapped to both this and other copies of the repeat (“Multi”). **D**, Microhomologies at insert breakpoints are often short (median, 1 bp). **(E–H)** Inverted insertions derived locally from the hotspot pair. **E**, Deletions with inverted insertions at the Chr19 hotspot pair. Inserts are derived in inverted orientation from the left or the right hotspot. For the Chr1 hotspot pair, see **Fig. S10B**. **F**, Example of an inverted insertion (top) and possible mechanisms (bottom). The right insertion breakpoint shares a 5-bp inverted microhomology with the left deletion breakpoint, while the left insertion breakpoint and the right deletion breakpoint do not share microhomology. Potential models explaining the inversion mechanism involve double cutting at the left hotspot or overhang foldback. **G**, Microhomologies at breakpoints for insertions derived from the Chr1 and Chr19 hotspot pairs. Inserts mapping to the left hotspot in a pair have long microhomologies present at their right breakpoints. Inserts mapping to the right hotspot mirror this pattern. See also **Table S5**. **H**, Inserts from the Chr1 and Chr19 hotspot pairs map close to hotspot centers.

Most of these inserts could be uniquely mapped, however, three were derived from the non-pseudoautosomal region (non-PAR) of the Y chromosome. The mouse non-PAR Y is highly ampliconic (Soh et al., 2014), so these inserts cannot be assigned to a single location. Hotspots have been called at some of these locations based on the assignment of unique SPO11 oligos, although most SPO11 oligos map to multiple locations that are highly similar (**Fig. 5B,C, S10A, S11**; **Table S4**). Among all chromosomes, the non-PAR Y experiences the largest increase in SPO11-oligo density in the absence of ATM (~30 fold) (Lange et al., 2016).

Inserts mapped at or close to the hotspot/cluster centers, such that nearly half had an end within 50 bp (**Fig. 5B**). We envision that these inserts derive from DNA fragments released by SPO11 double cutting; we infer that were not circularized because they are not permuted. It seems remarkable that inserts can sometimes originate from weak SPO11-oligo clusters, i.e., genomic sites that are much less frequently targeted by SPO11 than called hotspots, as such weak sites would be unlikely to experience two DSBs in the same cell if each DSB occurred independently at a population-average frequency. However, these findings are readily explained by the recent observation of concerted SPO11 cleavage within hotspots (especially in *Atm*^−/−^) (Johnson et al., 2019b). The ectopic insertion of SPO11-generated fragments into other SPO11-generated DSB locations represents a novel pathway for meiosis-specific, nonreciprocal genome modification by translocation of sequences.

#### Inverted insertions derived locally from the hotspot pair

Twelve deletions had insertions of 16 to 709 bp that mapped to one of the hotspots but were reinserted in opposite orientation (inverted insertions; **Fig. 5E, S10A,B**), including one with an inverted insertion from each hotspot in the pair. Reconstructing the insertion breaks illuminated potential inversion mechanisms. Specifically, in each case, one insertion breakpoint shared an inverted microhomology with one deletion breakpoint (median 5 bp) (**Fig. 5F,G**). The other insertion breakpoint, which was at or near this deletion breakpoint, had no or little microhomology with the other deletion breakpoint (median 0 bp). Inserts from the left and right hotspots mirrored each other in the sense that those from the left hotspot had the longer microhomology at the right breakpoint, whereas those from the right hotspot had the longer microhomology at the left breakpoint (**Fig. 5E,G, S10B**; **Table S5**). Insert sequences clustered around hotspot centers (**Fig. 5H**), indicating that they originated from DSB formation.

We propose two models to explain these events (**Fig. 5F**). In one, DSBs occur at or near the two inverted microhomologies, creating a small double-cut fragment within one hotspot, and a third DSB in the other hotspot generates a larger fragment destined for deletion. One end of the small fragment remains unprocessed, preventing normal recombination at this hotspot, while both ends with the inverted microhomology are at least partially resected, allowing strand annealing and fragment inversion. The unprocessed end of the now inverted fragment joins to the third DSB at the other hotspot. This is an attractive model because it incorporates defects seen or inferred in *Atm*^−/−^ mice, namely, impaired DSB processing (Paiano et al., 2020; Yamada et al., 2020) and closely spaced DSBs. In the second model, a DSB occurs next to just one of the inverted microhomologies and resection exposes the second microhomology. A foldback structure formed by annealing the inverted microhomologies is further processed by a structure-specific nuclease, generating a 5′ overhang that is filled in prior to joining the DSB end at the other hotspot. Foldback structures may be common in *Atm*^−/−^, because the excess of DSBs may exhaust the pool of RPA (Toledo et al., 2013), which normally prevents formation of secondary structures (Deng et al., 2015).

Interestingly, four inverted insertions from the right hotspot at the Chr1 hotspot pair involved the previously identified recurrent deletion site (**Fig. 4B, S10B**). In this case, 13 bp overlapping the recurrent deletion shares inverted microhomology with the most prominent DSB peak. Thus, this genomic location often engages in inappropriate repair (9% of all Chr1 deletions), demonstrating that DSB formation at or nearby such sequences may cause (mis)repair products to arise.

### Microdeletions within a single hotspot

If more than one DSB can form inside the same hotspot, and not just at adjacent hotspots, we predicted that small deletions might also arise within single hotspots. The presumed origin of insertions from hotspot fragments implies that closely spaced DSBs can indeed occur within individual hotspots. We applied deep sequencing to test for such microdeletions. We selected a hotspot on Chr1 ranked among the five strongest in both *Atm*^−/−^ and wild type. The SPO11-oligo profile for this hotspot displays a major central peak, a secondary peak, and several minor peaks, with SPO11 oligos spreading up to ~500 bp on both sides of the central peak, especially to the right (**Fig. 6A**). We designed four overlapping amplicons, ~580 bp each, to scan ~900 bp of the hotspot. This strategy would recover deletions around the major peak as well as those with one or both breakpoints away from the major peak, as often seen for inserts (**Fig. 6B**). Multiple, independent PCRs were carried out, each with input testis DNA equivalent to ~16,000 haploid genomes, for a total of ~10 million haploid genomes per amplicon. From pools of amplified DNA we sequenced ~3.5–5 million molecules per amplicon.

**Figure 6.**
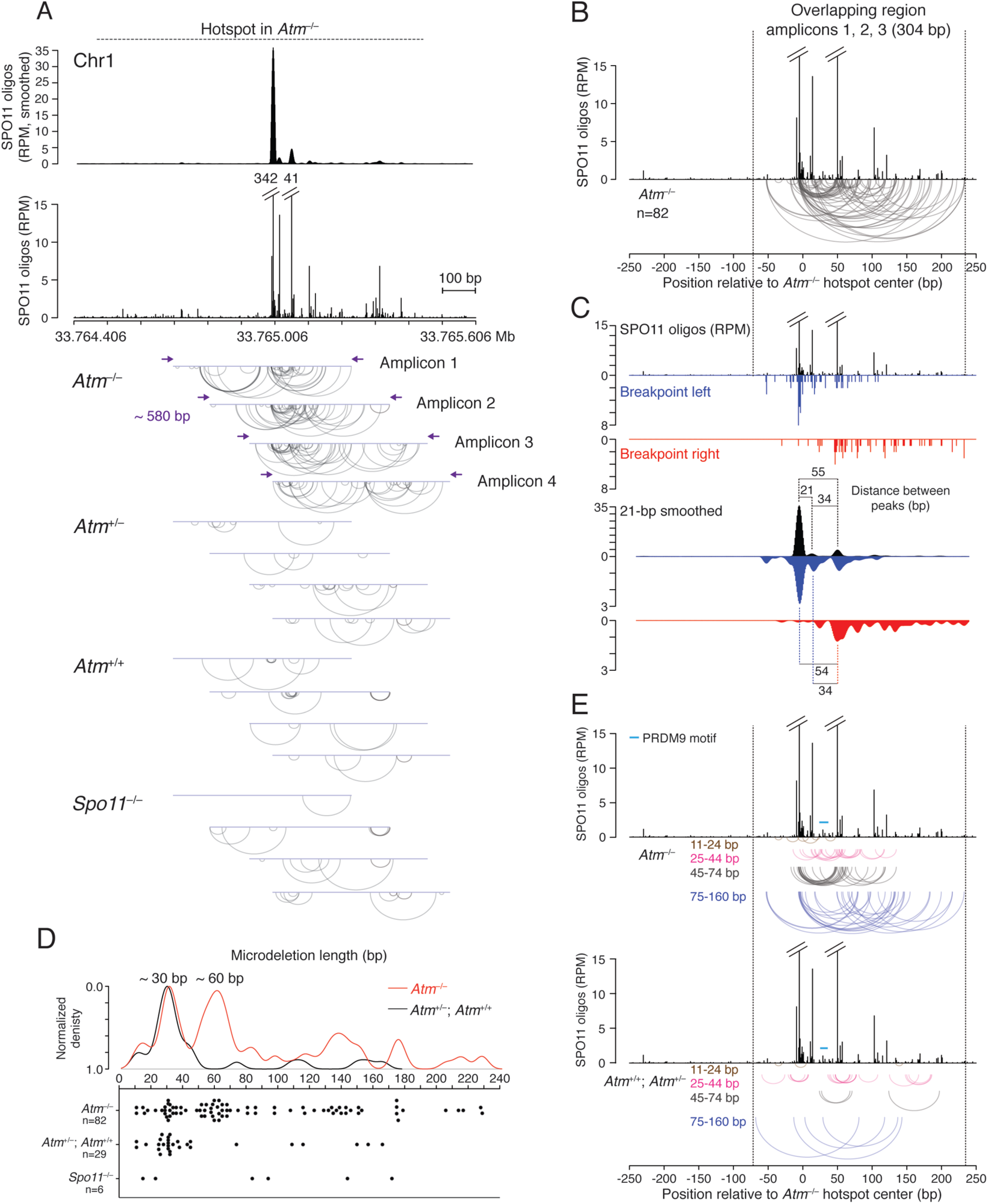
Microdeletions within a single DSB hotspot. **(A)** Detection of microdeletions at a strong DSB hotspot on Chr1 using amplicon deep sequencing. The hotspot displays a major SPO11-oligo peak, a secondary peak, and several minor peaks, as shown in the smoothed (21-bp Hann filter) and unsmoothed profiles. Four overlapping amplicons were deep sequenced to identify microdeletions. Arc diagrams of independent microdeletions found in each amplicon for the indicated genotypes are shown. For microdeletion frequencies, see **Table S6**. **(B, C)** Microdeletions from *Atm*^−/−^ in the region overlapping in amplicons 1, 2, and 3. **B**, Arc diagram of independent microdeletions. **C**, Distributions of the left and right breakpoints relative to the SPO11-oligo distribution. Both unsmoothed and smoothed profiles are shown. **(D, E)** Microdeletion length distributions in the overlapping region. **D,** Density and scatter plots of microdeletion lengths (>10 bp). **E,** Arc diagrams of microdeletions stratified by length. Microdeletions >160 bp are not shown, as their location may be more restricted by the size of the assessed region. See also **Fig. S13B,C** for larger microdeletion distribution.

We identified reads containing microdeletions ranging from 11 to 229 bp (**Fig. 6A**). (Reads with ≤10-bp deletions were not considered; **Fig. S12A**). In *Atm*^−/−^, 216 reads with microdeletions were detected from the four amplicons, for a total frequency of 1.58 × 10^-5^, significantly greater than in *Spo11*^−/−^ (0.19 × 10^-5^), wild type (0.54 × 10^-5^), and *Atm*^+/−^ (0.23 × 10^-5^) (p<0.0001, Chi square test) (**Table S5**). For some amplicons, reads from wild type and *Atm*^+/−^ were also more frequent than in *Spo11*^−/−^ (**Table S5**). If we disregard duplicate reads for a specific microdeletion from a given amplicon, we conservatively estimate that we obtained 149 independent microdeletions in total from *Atm*^−/−^ (**Fig. 6A**). This is likely an underestimate since some microdeletions were found in multiple amplicons (**Fig. S12B**).

Microdeletion breakpoints frequently clustered at SPO11-oligo sites in *Atm*^−/−^ (**Fig. 6A**). Prominent clusters were observed at the major SPO11-oligo peak (amplicons 1 to 3) and at the secondary SPO11-oligo peak (amplicons 1 to 4), indicating that microdeletions arose from SPO11-generated DSBs. This was even more evident when we pooled the 82 independent microdeletions from amplicons 1, 2, and 3 for which both breakpoints mapped within the 304-bp overlapping region (i.e., between the forward primer of amplicon 3 and the reverse primer of amplicon 1) (**Fig. 6B**). It was visually striking that many microdeletions had both breakpoints within these two SPO11-oligo peaks (17/82; 21%, 3 of which were recurring; **Fig. S12B**), implying that they originated from double cutting. In the smoothed breakpoint profiles, the most prominent left breakpoint peak matched the major SPO11-oligo peak, while the most prominent right breakpoint peak matched the secondary SPO11-oligo peak (**Fig. 6C**). These breakpoint peaks became even more prominent when all 116 reads in this overlap region were considered, something not seen with random sampling of reads (**Fig. S12C**). Thus, at least some of the duplicated reads likely represent independent microdeletions.

Microdeletions in the overlap region displayed a distinct length distribution (**Fig. 6D**), with two prominent microdeletion classes of ~30 bp and ~60 bp. The ~60-bp microdeletions predominantly mapped between the major SPO11-oligo peak and the secondary SPO11-oligo peak (71%, 17/24 from 49–70 bp; **Fig. 6E**), which are separated by 55 bp (**Fig. 6C**); reads within this class were often duplicated (**Fig. S13A**). A subset of the ~30-bp microdeletions (22%, 4/18 from 27–42 bp; **Fig. 6E**) mapped between a minor SPO11-oligo peak and the secondary SPO11-oligo peak 34 bp away (**Fig. 6C**). These results imply that hotspot substructure predicts double cutting and, hence, microdeletion locations. Conversely, double cutting will affect SPO11-oligo distribution within a hotspot when SPO11 oligos predominantly arise from double cutting.

In addition to microdeletions between two SPO11-oligo peaks, ~30-bp (56%, 10/18) and ~60-bp (21%, 5/24) microdeletions often had only one breakpoint at a SPO11-oligo peak (**Fig. 6E**). Assuming they also arose from two SPO11 breaks, the distinct size classes may reflect a more general underlying mechanism for double cutting, such that the length of double-cut fragments may be constrained. In that case, it remains possible that some of the ~60-bp microdeletions may have arisen from two double cuts (triple cutting), releasing two ~30-bp fragments, rather than a single double cut, although this may be less likely given that very few of the ~30-bp microdeletions have left breakpoints within the major SPO11-oligo peak like the ~60-bp microdeletions.

Microdeletions, especially the ~60-bp class, frequently removed the PRDM9 binding site, which is located between the major and secondary SPO11-oligo peaks (**Fig. 6E**). PRDM9 binding could potentially affect the spacing of double cuts. However, as similar microdeletion lengths are observed outside the PRDM9 motif, another or additional factors must exist that lead to this size class.

Interestingly, the major SPO11-oligo peak and the adjacent minor peak are separated by 21 bp (**Fig. 6C**), but the few small microdeletions (11–24 bp) did not map between these peaks (**Fig. 6D, E**), implying that two SPO11-induced DSBs may be constrained to longer distances. Thus, it is possible that they originated from single DSBs or even from double cuts with partial gap filling.

A less prominent microdeletion class of ~140 bp was also observed (129–153 bp, **Fig. 6D**). In several of these longer microdeletions, the left breakpoint clustered at the major SPO11-oligo peak, whereas the right breakpoint was located at sites in which fewer SPO11 oligos mapped (46%, 6/13; **Fig. 6E**). These longer microdeletions may also arise from double cutting. Long SPO11 oligos are observed in *Atm*^−/−^ testes (Lange et al., 2011), but are underrepresented in SPO11-oligo maps because of size selection during library preparation (Lange et al., 2016; Thacker et al., 2014). Even longer microdeletions were also observed that peaked at ~180 bp and ~230 bp (**Fig. 6D**), especially when considering all amplicons where microdeletions were less size-constrained (**Fig. S13B,C**). Some of these long microdeletions may have arisen from multiple double-cut fragments that generated long gaps. However, it is notable that the median insert size from other hotspots in the paired hotspot deletions was quite long (~100 bp), with one insert extending 349 bp, suggesting that long double-cut fragments can be generated.

In *Atm*^+/+^ and *Atm*^+/−^, microdeletions did not display obvious spatial patterns (**Fig. 6A,E**). Because the frequency of microdeletions in the presence of ATM is only slightly higher in some amplicons than in *Spo11*^−/−^, we consider that a portion of these events arose from PCR errors or other genomic damage, as in *Spo11*^−/−^. Strikingly, however, a ~30-bp (25–43-bp) microdeletion class was also apparent, but not in the *Spo11*^−/−^ control, although the prominent ~60-bp microdeletion class found in *Atm*^−/−^ was not observed (**Fig. 6D, S13B**). Microdeletions from the ~30-bp class did not map within SPO11-oligo peaks, although some breakpoints clustered (**Fig. 6E**). Possibly, in the presence of ATM, microdeletions are more likely to occur at DSBs formed outside of SPO11-oligo peaks. In both the presence and absence of ATM, microhomology at breakpoints was generally short (median 1 bp) (**Fig. S13D**), similar to deletions between hotspot pairs.

Our assumption that SPO11 double cutting occurs within individual hotspots, preceding microdeletion formation, is consistent with recent findings from *S. cerevisiae* in which double-cut fragments were observed ((Johnson et al., 2019b); Franz Klein, personal communication). Their minimum size was determined to be ~30 bp but longer fragments in increments of ~10 bp were abundant as well. The authors suggest models in which Spo11 is constrained to make the simultaneous cleavages. The recent finding of condensates of the yeast DSB repair machinery suggests a mechanism whereby centers of DSB activity form which could provide such a platform (Claeys Bouuaert et al., 2020). Our single hotspot data fit well with both of these concepts. Further, potential SPO11 double-cut fragments have been computationally identified within mouse hotspots (Johnson et al., 2019b) from SPO11-oligo libraries (Lange et al., 2016). Assuming a similar lower size limit in mouse as in yeast, double-cut fragments are predicted to peak at ~32 bp and extend to ≤60 bp in *Atm*^−/−^, near the size limit of the SPO11-oligo library preparation (Lange et al., 2016). Our analysis of microdeletions is in line with this finding, given a prominent microdeletion class of ~30 bp, but also suggests that SPO11-oligo double-cut fragments extend much longer, i.e., from ≥60 bp to at least ~230 bp, congruent with the observation of long SPO11 oligos in *Atm*^−/−^ (Lange et al., 2011).

## Conclusions

Our study uncovers an unexpected pathway for germline mutagenesis in mice involving nonhomologous end-joining at SPO11 double cuts and leading to various mutational outcomes. Although pronounced in the absence of ATM, these *de novo* mutations also occur at low frequency in the presence of ATM. Deletions and tandem duplications result in copy number variations (CNVs), predicting a role in genome evolution (Collins et al., 2020). Moreover, deletions and duplications in both our hotspot pairs remove and duplicate exons, respectively, demonstrating that SPO11-mediated events can disrupt or otherwise modify gene function. The occurrence of events involving two chromosomes also predicts gross rearrangements. Thus, our findings reveal a previously hidden potential in meiosis for mutagenesis and the disruption of genome integrity. Furthermore, double-cut fragments appear to be released from hotspots and inserted at other genomic sites undergoing DSBs, also with mutagenic consequences. Because DNA gaps are thought to initiate a subset of meiotic recombination events as part of the normal meiotic program (Johnson et al., 2019b), this raises the question as to how re-insertion of released double-cut fragments is usually suppressed. We speculate that circularization of these fragments could be a mechanism to suppress reintegration into the genome.

Microdeletions resulted in PRDM9 motif deletion in 38% of events in the absence of ATM and 11% in the presence of ATM. By eliminating PRDM9 motifs, such deletions shape the recombination landscape, thus influencing genome evolution (Baker et al., 2015; Cole et al., 2014) and potentially contributing to speciation (Davies et al., 2016; Smagulova et al., 2016). Because PRDM9 drives recombination away from promoters (Brick et al., 2012), deletions may be better tolerated in PRDM9-dependent meiotic recombination, including in humans, and in turn have a greater potential impact on human genome evolution.

## Acknowledgments

We thank Matthew Neale and Franz Klein for sharing unpublished results; members of the Jasin and Keeney labs for assistance and discussions, in particular Shintaro Yamada; and MSK core facilities, in particular the Integrated Genomics Operation (Cassidy Cobbs, Neeman Mohibullah, Agnès Viale) and the Bioinformatics Core (Nicholas Socci), which are supported by NIH grant P30 CA008748. This work was supported by March of Dimes grant 1-FY17-799 (A.L.), American Cancer Society fellowship PF-12-157-01-DMC (J.L.), NIH grant R35 GM118092 (S.K.), and NIH grant R35 GM118175 (M.J.).

## Author contributions

A.L., S.K., and M.J. conceived the research and designed the experiments with contributions from J.L.

A.L. performed all of the experiments. A.L. and M.J. wrote the manuscript with input from S.K. and J.L.

## Methods

### Mouse husbandry

All experiments were conducted according to US Office of Laboratory Animal Welfare regulatory standards and were approved by the Memorial Sloan Kettering Cancer Center Institutional Animal Care and Use Committee. Mice were purchased from The Jackson Laboratory where indicated. *Atm*^+/−^ males on the C57BL/6J (B6) background (B6.129S6-*Atm^tm1Awb^*/J; stock #008536) were crossed with B6 females (stock #000664) to establish a breeding colony and to obtain experimental animals. The *Atm* mutation was previously described (Barlow et al., 1996). *Atm*^+/−^ B6 mice were crossed with *Atm*^+/−^ DBA/2J (DBA) mice to get F1 hybrid males. *Atm*^+/−^ mice on the DBA background were obtained by six backcrosses of *Atm*^+/−^ mice that were partially on an A/J background with DBA inbred mice (stock #000671). Animals were fed regular rodent chow with continuous access to food and water.

### Isolation of testis and sperm DNA

DNA was isolated from testicular single-cell suspensions to enrich for germ cells. Testis dissociation was based on protocols previously described (Cole et al., 2014; Gaysinskaya et al., 2014) with some modifications. Testes from juvenile 15-day-old and adult 2- to 5-month-old mice were decapsulated and seminiferous tubules incubated at 33 °C in 10 ml Gey’s Balanced Salt Solution (GBSS) (Sigma) with 0.5 mg/ml collagenase (Sigma) and 2 μg/ml DNase I (Roche) for 15–20 min, shaking at 500 rpm. After 10 min and at the end of the incubation the tubules were gently pipetted up and down using a transfer pipette. The tubules were washed twice with 10 ml GBSS and digested with 0.5 mg/ml trypsin (Sigma) in 10 ml GBSS, supplemented with 2 μg/ml DNase I, for 25–40 min at 33 °C, shaking at 500 rpm. Pipetting up and down was performed at the start, midpoint and end of the incubation. Trypsin was inhibited by adding 10% fetal calf serum. Cell suspension was obtained by repeated pipetting for 2 min and filtration through a 70-μm cell strainer (ThermoFisher). All of the mutant cell suspension and one-third to one-half of the wild-type cell suspension was transferred to a 15 ml tube and centrifuged for 3 min at 1550 × g. The cell pellet was resuspended by tapping the tube, followed by the addition of 600 μl lysis buffer (200 mM NaCl, 100 mM Tris-Cl pH 7.5, 5 mM EDTA, 0.5% SDS, 0.5 mg/ml Proteinase K). The lysate was transferred to a 1.5 ml Eppendorf tube and incubated overnight at 55 °C. Genomic DNA was isolated using phenol-chloroform extraction and alcohol precipitation, with the extraction step repeated twice. DNA was dissolved in 200–400 ml H_2_O, incubated for 1 h at 55 °C and overnight at 4 °C. DNA concentration was quantified by NanoDrop (ThermoFisher). DNA was diluted to ~200 ng/μl and incubated for another 2–3 h at 37 °C to reduce the viscosity of the DNA solution. DNA concentration was measured again three times before freezing. DNA samples were stored at −20 °C.

Sperm DNA was isolated as previously described (Peterson et al., 2020).

### Detection of deletions at hotspot pairs by nested PCR

Deletions were detected by nested PCR using primers flanking the two hotspots of a pair (**Fig. 1B, S2A**). Prior to PCR, genomic DNA was digested with restriction enzymes where indicated to suppress parental DNA amplification (**Fig. S4**). Restriction digestions were performed in a 60 μl volume with 1200 ng of DNA and 4–4.5 μl restriction enzyme(s) (Chr1 hotspot pair: NsiI, NcoI, and BglII or NcoI alone; Chr19 hotspot pair: BamHI or KpnI; BioLabs). Reactions were incubated overnight at 37 °C followed by an additional 3 h of incubation with a 1.5–2 μl of fresh restriction enzyme(s), then placed on ice before PCR. In the majority of PCRs, 2.5 μl of the digestion reaction was used per PCR, equivalent to 50 ng of genomic DNA, with this DNA concentration being optimal for the estimation of deletion frequencies; 1.25 ml and 5 ml, equivalent to 25 ng and 100 ng, respectively, were also tested. In the experiments with no digestion, 50 ng of DNA template was used per PCR in most cases.

Nested PCR was based on the previously described protocol (Cole and Jasin, 2011) with some adjustments. A first round of PCR was performed using the following primers: Chr1_DH_FW1: 5′-GGTCAGGAGGATGGGGAG-3′ and Chr1_DH_RV1: 5′-CCAAGAAGGTGCTTATCGTCC-3′ for the Chr1 hotspot pair, Chr19_DH_FW2: 5′-TGGAATGTAGCCTCTGGCAG-3′ and Chr19_DH_RV2: 5′-GTGCTGTCAGAAGCCGTGG-3′ for the Chr19 hotspot pair, Chr1_DH_DB_FW1: 5′-CCTGGAGACCAAAGCCAC-3′ and Chr1_DH_RV1: 5′-CCAAGAAGGTGCTTATCGTCC-3′ for the Chr1 hotspot pair in B6/DBA F1 hybrids. A second round of PCR was performed using the following primers: Chr1_DH_FW2: 5′-GCCTTGGCTCAAAGTGTCTG-3′ and Chr1_DH_RV2: 5′-TCCCTTCTGCCATCCTTCTG-3′ for the Chr1 hotspot pair, newChr19_DH_FW2: 5′-CCCAAAGTTGTCCTTGCTTCC-3′ and newChr19_DH_RV2: 5′-TGGTTTCCAGAGGAGAGGAC-3′ for the Chr19 hotspot pair, Chr1_DH_DB_FW2: 5′-GGCATTGCTTCATTCATGGTCTG-3′ and Chr1_DH_RV2-3: 5′-CATTGTCATCTTGAGCACTCCTC-3′ for the Chr1 hotspot pair in B6/DBA F1 hybrids. The annealing temperature was optimized for each primer pair by gradient PCR with testis genomic DNA of known high quality. DNA samples from different testes were also tested for the appearance of a strong parental PCR band using the primary PCR primers.

For nested PCR, each primary PCR was performed in a 50 μl volume containing: 1x Jeffreys buffer (10x: 450 mM Tris-Cl pH 8.8, 110 mM (NH_4_)_2_SO_4_, 45 mM MgCl_2_, 67 mM 2-mercaptoethanol, 44 μM EDTA, 10 mM each dNTP (Roche), 1.13 mg/ml ultra-pure (non-acetylated) BSA (Ambion), 12.5 mM Tris base), 0.2 μM primers, 0.03 U/μl Taq DNA polymerase (ThermoFisher) and 0.006 U/μl Pfu Turbo DNA polymerase (Agilent). The primary PCR product (0.6 μl) was treated with S1 nuclease (ThermoFisher) at room temperature for 30 min, in a 5 μl volume containing: 0.5x S1 buffer, 0.2 U/μl S1 nuclease and 5 ng/μl sonicated salmon sperm DNA. The reaction was diluted with 45 μl of dilution buffer (10 mM Tris-Cl pH 7.5 and 5 ng/μl sonicated salmon sperm DNA). 4 μl of the diluted S1 reaction was seeded into the secondary PCR, which was performed in a 20 μl volume containing the same components as the primary PCR. Deletion products were identified by electrophoresis on a 1% agarose gel (**Fig. 1C, S2B**). PCR products smaller than the parental size were excised, purified, and Sanger sequenced using primers from the second round of PCR and in some cases additional primers. Diluted S1 reactions were stored at −20 °C, and used again for secondary PCRs if more PCR product was needed for sequencing.

PCR conditions were tested for each hotspot pair. The best result, that is, the appearance of more intense, smaller PCR products, was obtained when the extension time was shortened (<60 s/kb) in the primary PCR to inhibit amplification of the longer parental PCR product. The extension time in the secondary PCR was slightly shorter.

Primary PCR conditions were denaturation at 96 °C for 2 min, followed by 37 cycles of amplification as follows: denaturation at 96 °C for 20 s, annealing at 59 °C for 30 s, extension at 65 °C for 2 min for the Chr1 and Chr19 hotspot pairs (3.6 kb parental products), and 2 min 30 s for the Chr1 hotspot pair in B6/DBA hybrids (4 kb parental product). Secondary PCR conditions where the same, but with different extension times: 3 min for the Chr1 and Chr19 hotspot pairs (3.4-kb parental products) and 3 min 30 s for the Chr1 hotspot pair in B6/DBA hybrids (3.9-kb parental products).

Sequenced PCR products were analyzed using ApE software (https://jorgensen.biology.utah.edu/wayned/ape/) to identify deletions, microhomologies, and polymorphisms in DNA sequences. Deletion breakpoint profiling was performed using R versions 3.4.4 to 3.6.3 (http://www.r-project.org). Graphing was performed in R and GraphPad Prism version 8.

### Detection of tandem duplications and circular DNA by nested inverse PCR

Primary and secondary PCR reaction protocols, including S1 digest, were the same as in the nested PCR protocol detecting deletions. For the primary PCR, the following primers were used: Chr1_Inv_FW1: 5′-CACTCATTCCCACTTGGCTC-3′ and Chr1_Inv_RV4: 5′-GGACGCATGATAAGCAAGGAG-3′ for the Chr1 hotspot pair, Chr19_Inv_FW3: 5′-TTCTGAGAGAGTCTTGCATTGG-3′ and Chr19_Inv_RV1: 5′-CCTGTACTGTTCATTCGCCATC-3′ for the Chr19 hotspot pair. For the secondary PCR, the following primers were used: Chr1_Inv_FW2: 5′-CAATCCGATGGCATTTAGATTAGG-3′ and Chr1_Inv_RV5: 5′-GGAGAAATGAGTTAATGAAATAGCC-3′ for the Chr1 hotspot pair, newChr19_Inv_FW4: 5′-CTCAGTGCTGGAATTACAAGAG-3′ and Chr19_Inv_RV2: 5′-CACTAAACCGGAGTTCGCTG-3′ for the Chr19 hotspot pair. Inverse PCR does not amplify parental DNA. Therefore, we paired each primer with another primer to test for the appearance of the parental PCR product and to optimize the annealing temperature by gradient PCR. These control primers were the same primers we used to detect deletions.

Primary and secondary inverse PCR conditions were the same: 96 °C for 2 min followed by 30 cycles at 96 °C for 20 s, 61 °C for 30 s and 65 °C for 3–4 min. For both hotspot pairs, if double cutting occurs at hotspot centers, the expected PCR product is 1.4 kb and 1.3 kb for the primary and secondary PCR, respectively. In our experiments, the median size of secondary PCR products was 702 bp for Chr1 and 844 bp for Chr19. However, the more frequent appearance of smaller PCR products was not due to the extension time.

Inverse PCR product sequencing and further analysis were the same as for deletions.

### ExoV digestion prior to inverse PCR to detect circular DNA

ExoV was used to degrade linear DNA but not circular DNA. To ensure DNA degradation at the Chr1 hotspot pair we digested genomic DNA with the ScaI restriction enzyme (**Fig. 3D, S6B**). Restriction digestions were performed overnight at 37 °C in a 60 μl volume with 1200 ng of DNA, 4 μl ScaI (BioLabs), followed by 3 h of incubation with a fresh portion, 2 μl, of the enzyme. Digestions were inactivated at 80 °C for 20 min, and placed on ice. Next, reactions were split into two and half was treated with ExoV in a 60 μl final volume containing: 1x CutSmart buffer, 1mM ATP and 0.2 U/μl ExoV (or H_2_O). Reactions were incubated at 37 °C for 1–2 h and inactivated at 70 °C for 30 min, then placed on ice. To remove short oligonucleotides generated by ExoV, 3 μl of Thermolabile ExoI (BioLabs) was added to each reaction and samples were incubated for 15 min at 37 °C, 5 min at 80 °C, then placed on ice.

For each inverse PCR, 2.5 μl of the digestion reaction (+/– ExoV) was used, equivalent to 25 ng of pre-digestion genomic DNA. In parallel, a control PCR was performed to test ExoV efficiency. To this end, an aliquot of the digestion reaction was diluted (1:10, 1:100, and 1:1000), and used in nested PCR with control primers (reverse inverse primers plus forward primers used to detect deletions in B6 × DBA F1 hybrids), amplifying a 2.1-kb PCR product (**Fig. S6B**). PCR conditions for inverse PCR and control PCR were the same as described above.

### Detection of microdeletions at a single hotspot by amplicon deep sequencing

Four ~580-bp overlapping amplicons of the hotspot (**Fig. 6A**) were amplified using the following primer pairs: amplicon 1, Chr1_MiSeq_FW1: 5¢-TCATGTCCGGTTTGGATGCC-3¢ and Chr1_MiSeq_RV1: 5¢-CGAATCAAAGACTCCAGAAGAGG-3¢; amplicon 2, Chr1_MiSeq_FW2: 5¢-AGTAGCAAGAAGGGCTGGAG-3¢ and Chr1_MiSeq_RV2: 5¢-AGAGACAGATGTGGAGGCTAAC-3¢; amplicon 3, Chr1_MiSeq_FW4: 5¢-CTGAATCTCTCCTGGTGTGTC-3¢ and Chr1_MiSeq_RV4: 5¢-AGGACATTGAATGTTTTTATGGTTC-3¢; amplicon 4, Chr1_MiSeq_FW5: 5¢-AAACAGAAGAAGCAGTAATGCC-3¢ and Chr1_MiSeq_RV5: 5¢-GTGTGCAACTGGGTATATTTCAG-3¢. For each primer pair the annealing temperature was tested by gradient PCR with testis genomic DNA of known high quality. To minimize PCR bias and jackpot amplification of single deletion events, the number of amplification cycles was determined by when a faint PCR product was first visible, which was 22 cycles for all samples. For each amplicon, PCR conditions were the same: 96 °C for 2 min followed by 22 cycles at 96 °C for 20 s, 59 °C for 30 s and 65 °C for 35 s. PCR was performed in a 50 μl total containing 1.5 U Taq DNA polymerase (ThermoFisher) and 0.3 U Pfu Turbo DNA polymerase (Agilent) in the Jeffreys buffer described above for nested PCR with 50 ng of testis genomic DNA. Testis DNA was derived from 3-5 different mice for each genotype and amplified independently.

PCRs were carried out in 96-well plates to generate a large number of independently-derived PCR products for sequencing. For each amplicon, 576 PCRs were performed with a total of 28.8 μg genomic DNA, equivalent to 9.6 million haploid genomes. PCRs were pooled, and a 1200-μl aliquot was purified and concentrated on spin columns (Invitrogen PureLink^TM^ PCR Purification Kit) as follows: For each 100 μl PCR aliquot, 400 μl binding buffer from the kit was added, and then the sample was applied to the column and centrifuged. This step was repeated 6 times in each of two columns before washing and elution with 37 μl elution buffer. Eluted DNA was combined from the two columns and a 15-μl aliquot was run on a 2% agarose gel to estimate DNA concentration, which was ~2 ng/μl. The remaining ~60 μl was submitted to the Integrated Genomics Operation at MSK for library preparation and deep sequencing. The library was prepared by ligation of indexed adapter sequences using the KAPA HyperPrep Kit (Roche). Sequencing was performed using Illumina MiSeq (2 x 300-bp paired-end reads). In a single run, libraries from amplicons 1–4 were mixed with 5% PhiX DNA control library (Illumina) and sequenced. Because MiSeq can deliver 15–20 million reads per lane, in our experiment, 3.6–4.8 million reads per amplicon were estimated, which was about 2 times lower than the 9.6 million input DNA templates. With that calculation, not all deletion events were recovered, but most recovered deletions should have yielded only a single read.

We performed 4 MiSeq runs with amplicons from different genotypes (**Fig. S12A**). FASTQ sequencing files were analyzed using CRISPResso2 (Clement et al., 2019; Pinello et al., 2016) by the Bioinformatics Core Facility at MSK. Reads were aligned to the reference amplicon sequence and 2.9–5.3 million paired-end reads were mapped per amplicon. Alignments matching the target sequence for the first 20 and last 20 bases and with one deletion event were filtered. Reads with >1 bp deletion and with no mutations in the sequence were analyzed in R and ApE. Breakpoint distribution analyses and random sampling (**Fig. 12C**) were performed in R. Graphs were plotted using R and GraphPad Prism, and the Chi-square test was performed in GraphPad Prism.

### SPO11-oligo maps

SPO11-oligo maps were generated previously (GEO accession number GSE84689) (Lange et al., 2016). Mice for the *Atm*^−/−^ data set “*Atm* null 1”, which was used in all the analyses, were on a mixed B6 and 129/Sv strain background. Two *Atm*^+/+^ data sets are available, “B6”, from mice on a pure B6 background, and “*Atm* wt”, from the same colony as *Atm*^−/−^ mice. Hotspots selected in this study were called in the “B6” data set (**Fig. 1B, S1D, S2A**), while analysis of hotspot distributions was done with the “*Atm* wt” data set (**Fig. 1A, S1A–C**). The “B6” data set was also used to check if SPO11 oligos map to the genomic locations from which inserts derived (**Fig. S9, S11**).

### Data and code availability

Original data including gel pictures, Sanger sequencing, all data processing spreadsheets and R scripts are available from the lead contact upon request. Raw data from amplicon deep sequencing will be deposited in the GEO database.

---

**Figure S1.**
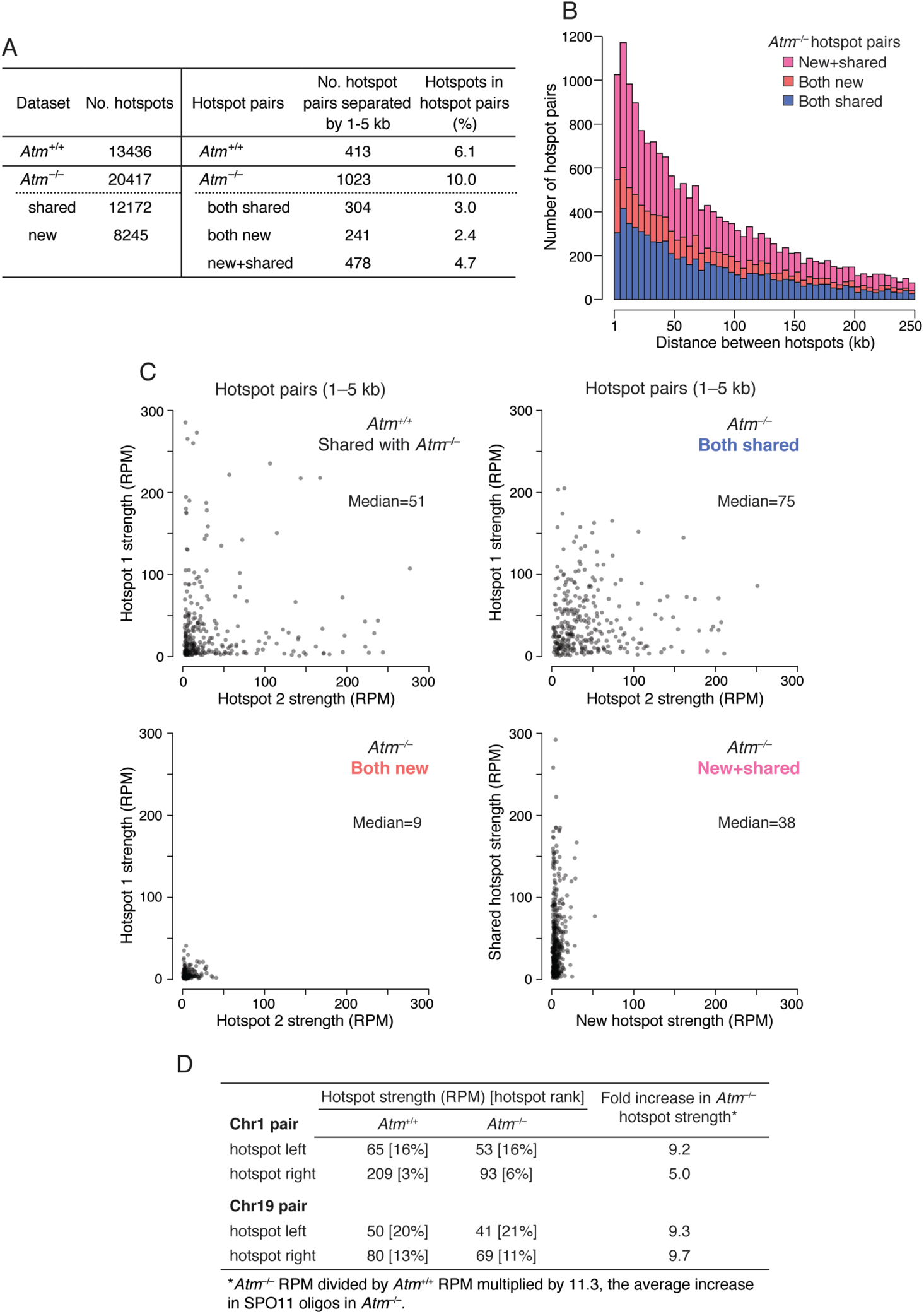
Higher potential in *Atm*^−/−^ for DSB formation at adjacent hotspots. Analyses were performed for autosomes. **(A)** Hotspot pairs separated by 1–5 kb are observed more frequently in *Atm*^−/−^ than in *Atm*^+/+^. Data are derived from SPO11-oligo maps (Lange et al., 2016). “Shared” hotspots are those called in both *Atm*^−/−^ and *Atm*^+/+^. “New” hotspots are those called only in *Atm*^−/−^, although corresponding SPO11-oligo clusters are typically present in the *Atm*^+/+^ map (Lange et al., 2016). Hotspot pairs in *Atm*^−/−^ are separated into classes according to whether each hotspot is shared or new. **(B)** Distribution of distances separating hotspots in *Atm*^−/−^ hotspot pairs, shown up to 250 kb. The proportion of each hotspot pair class from panel **A** is indicated. The increase in the number of hotspot pairs in *Atm*^−/−^ separated by short distances (Fig. 1A) is primarily the result of the emergence of new hotspots close to shared hotspots. **(C)** Strength of individual hotspots in hotspot pairs separated by 1–5 kb. Hotspot pair strength, the median of total RPM (reads per million) for the two hotspots in a pair, is indicated. Note that the RPM values do not account for the 11.3-fold increase in SPO11 oligos in *Atm*^−/−^ (Lange et al., 2011). Outliers with >300 RPM (4 in *Atm*^−/−^; 18 in *Atm*^+/+^) are not shown. **(D)** Hotspot strength for the individual hotspots in the Chr1 and Chr19 hotspot pairs. RPM values in *Atm*^−/−^ are multiplied by 11.3, the average increase in whole-testis SPO11-oligo levels in *Atm*^−/−^ mice. Hotspot rank is the rank of the individual hotspot within all hotspots for *Atm*^+/+^ or *Atm*^−/−^.

**Figure S2.**
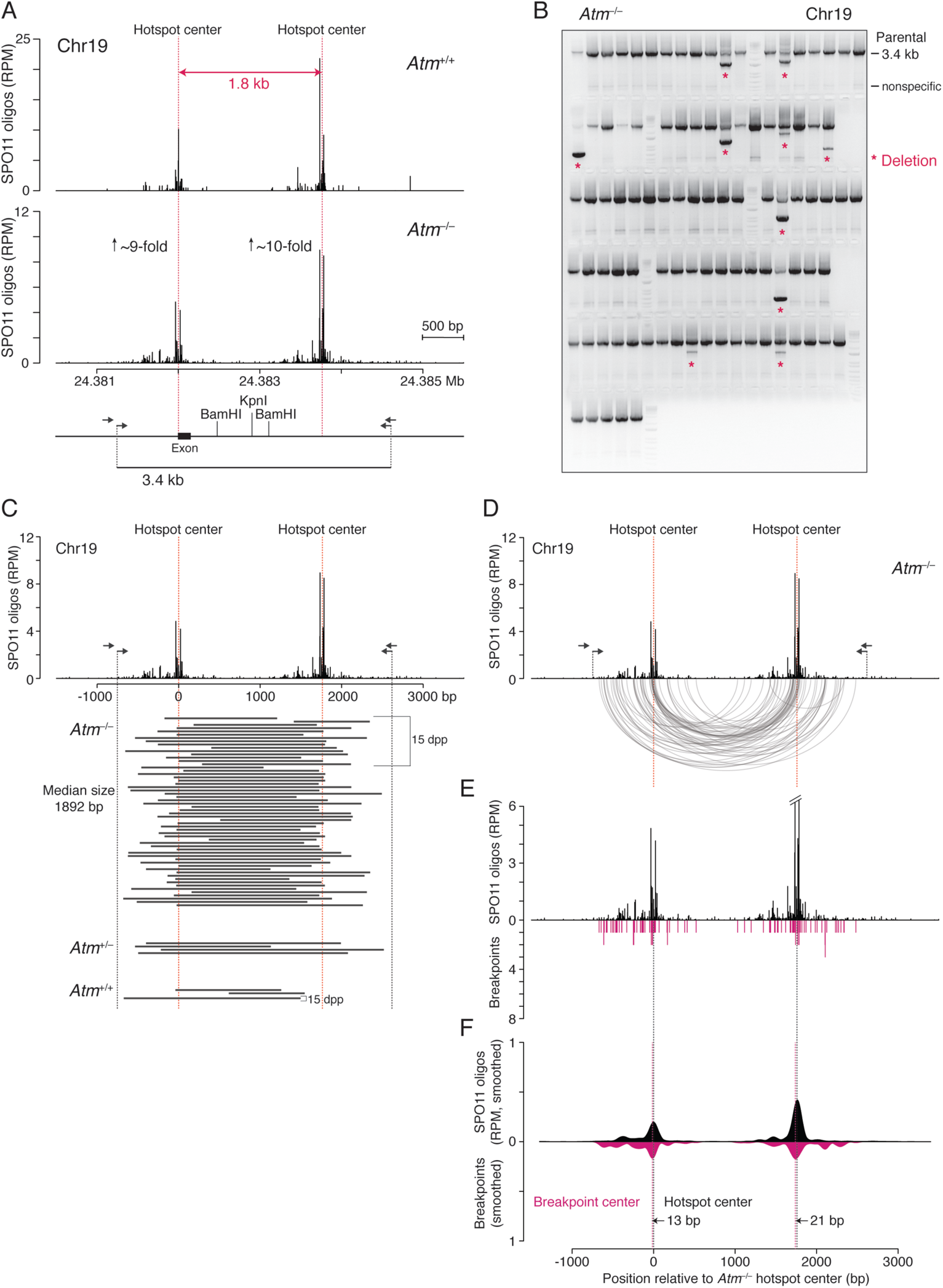
Deletions at the Chr19 hotspot pair. **(A)** SPO11-oligo maps in wild type and *Atm*^−/−^ at the Chr19 hotspot pair, with the scheme of the nested PCR assay below. The fold increase in the SPO11 oligos (RPM × 11.3) is indicated for each hotspot. Prior to amplification, genomic DNA is optionally digested with restriction enzymes to reduce amplification of parental products. The *Pip5k1b* exon located at the left hotspot is removed in many of the deletions. **(B)** Representative gel of PCR products in *Atm*^−/−^ from a 96-well plate. ~16,000 haploid genomes were seeded per PCR. Asterisks indicate deletions. **(C–F)** Analysis of deletions. **C**, Deleted DNA sequences (black bars) from adult and juvenile testes. **D**, Arc diagram of deletions in *Atm*^−/−^ (58 deletions total). See also **Fig. S3**. **E**, Distribution of individual breakpoints. **F**, Smoothed breakpoint and SPO11-oligo profiles (201-bp Hann filter for smoothing). Offsets between centers of hotspots and breakpoint distributions are indicated.

**Figure S3.**
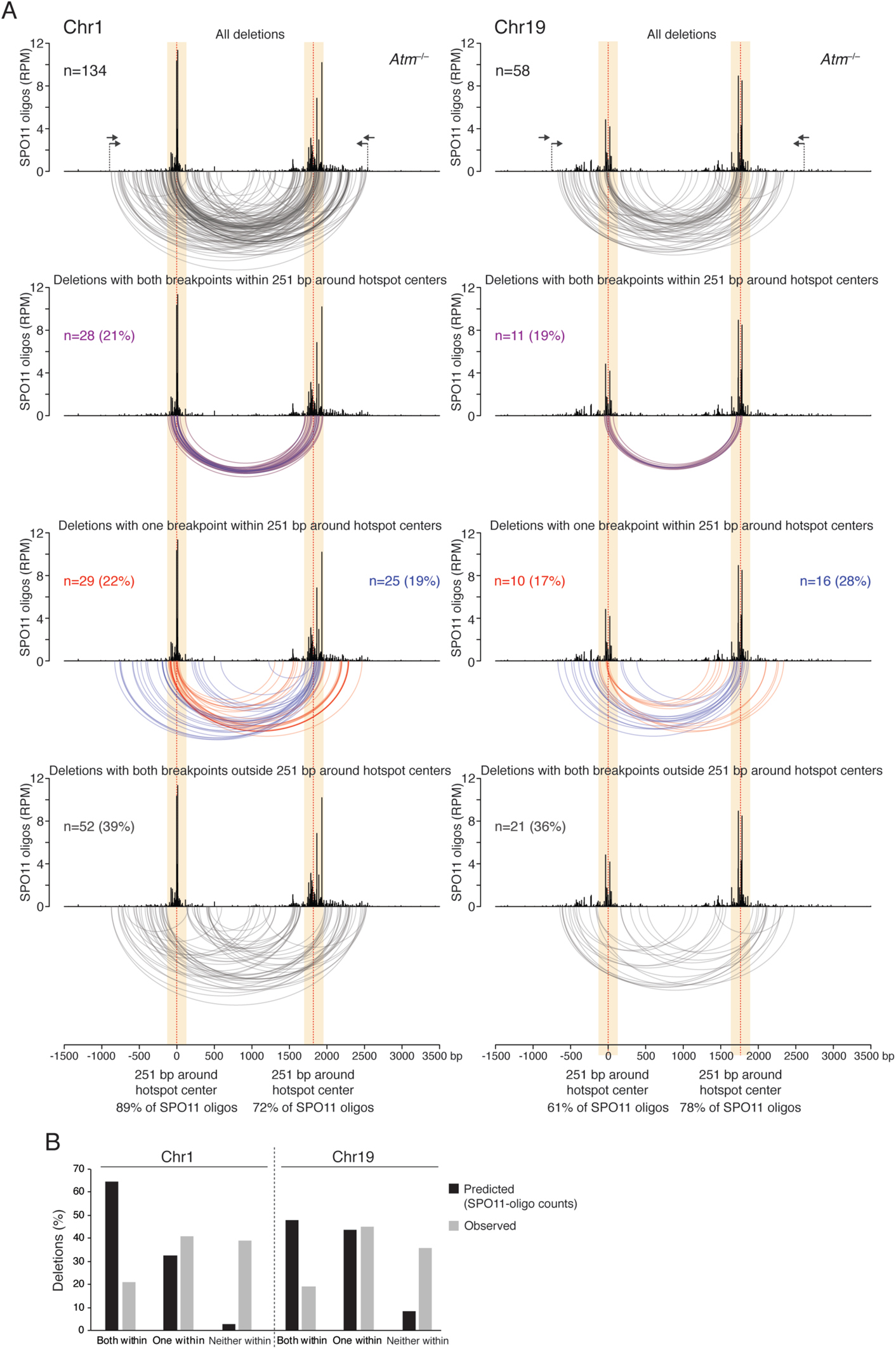
Distribution of deletions for the Chr1 and Chr19 hotspot pairs. **(A)** Deletion arcs from *Atm*^−/−^ are segregated according to the positions of breakpoints relative to 251-bp zones around each hotspot center (yellow). The percentage of SPO11 oligos is the number of SPO11 oligos within each 251-bp zone divided by the total number of SPO11 oligos mapping between the left or right primer (arrows) and the midpoint between hotspot centers. Deletions were grouped according to whether both breakpoints (violet arcs), one breakpoint (red and blue arcs), or neither breakpoint (gray arcs) mapped within the 251-bp zones. The percentage of deletions within each group is indicated. **(B)** Predicted and observed percentage of deletions with both breakpoints, one breakpoint, or neither breakpoint within 251 bp around hotspot centers. The predicted distribution of deletions is equivalent to the expected distribution of double cuts calculated from SPO11-oligo counts (RPMs) at the hotspot pairs.

**Figure S4.**
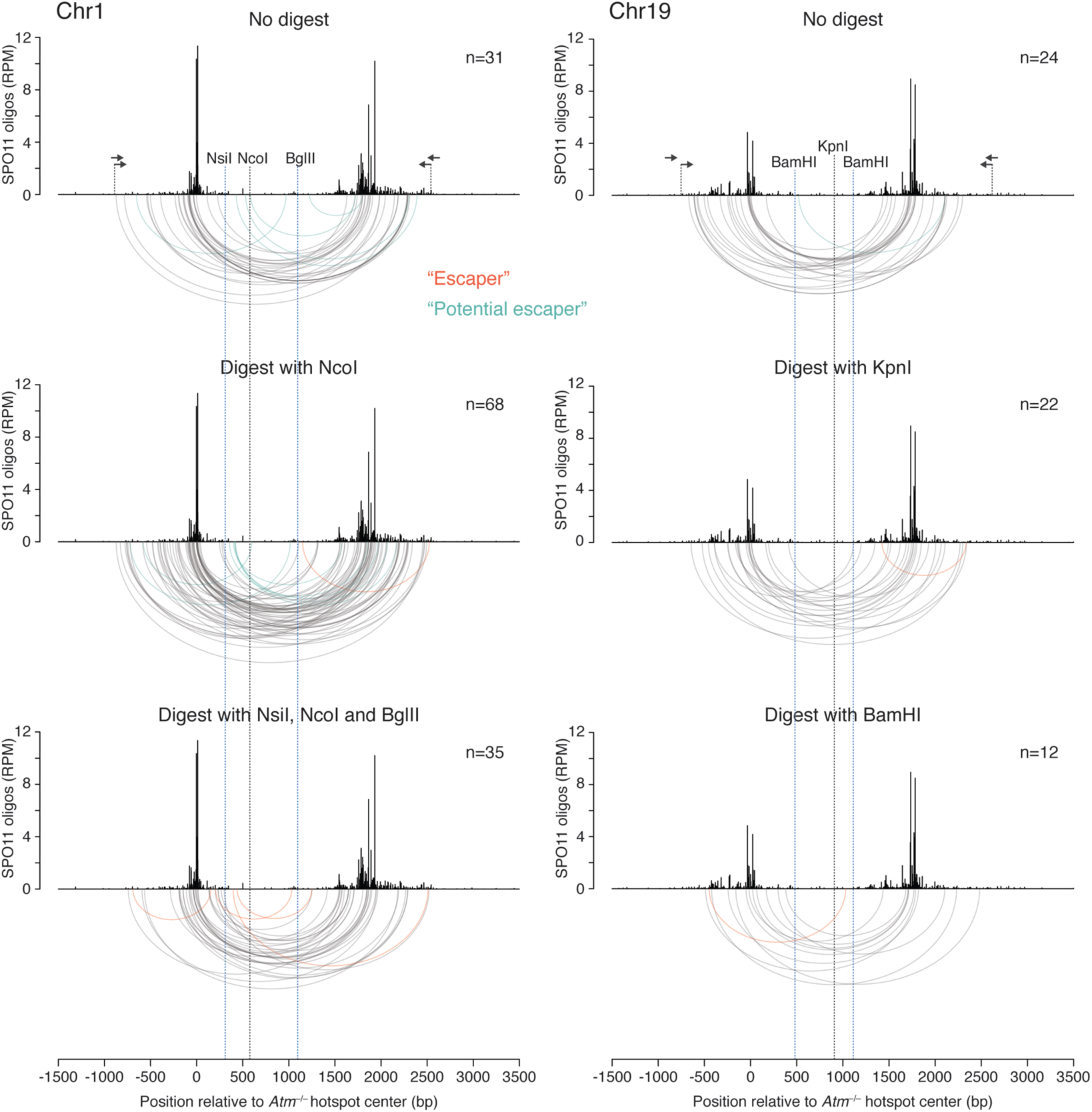
Distribution of deletions from experiments with or without prior genomic DNA digest. Genomic DNA was optionally digested with restriction enzymes prior to PCR to suppress amplification of parental DNA. Nested primers are indicated with arrows. Restriction sites located in between the two hotspots of a pair are expected to be absent in deletion products. Consistent with this, the majority of deletions in *Atm*^−/−^ at the Chr1 and Chr19 hotspot pairs had both breakpoints outside of the restriction site region (vertical blue lines). Similar results were obtained in experiments in which the digestion step was omitted, or only the enzyme cutting uniquely inside this region was used (NcoI or KpnI, vertical black lines). A few deletion products contained restriction sites due to incomplete genomic DNA digestion (“Escapers”, orange arcs) or were captured in experiments with no digest or when the unique central cutting enzyme was used (“Potential escapers”, green arcs). Overall, deletion distributions were similar in all experiments, independent of the digest, with breakpoints clustered around hotspot centers. Note that some of these deletions had both breakpoints mapping to one hotspot of the hotspot pair (4 at the Chr1 pair; 1 at the Chr19 pair), likely arising from DSBs formed within one hotspot, as with microdeletions.

**Figure S5.**
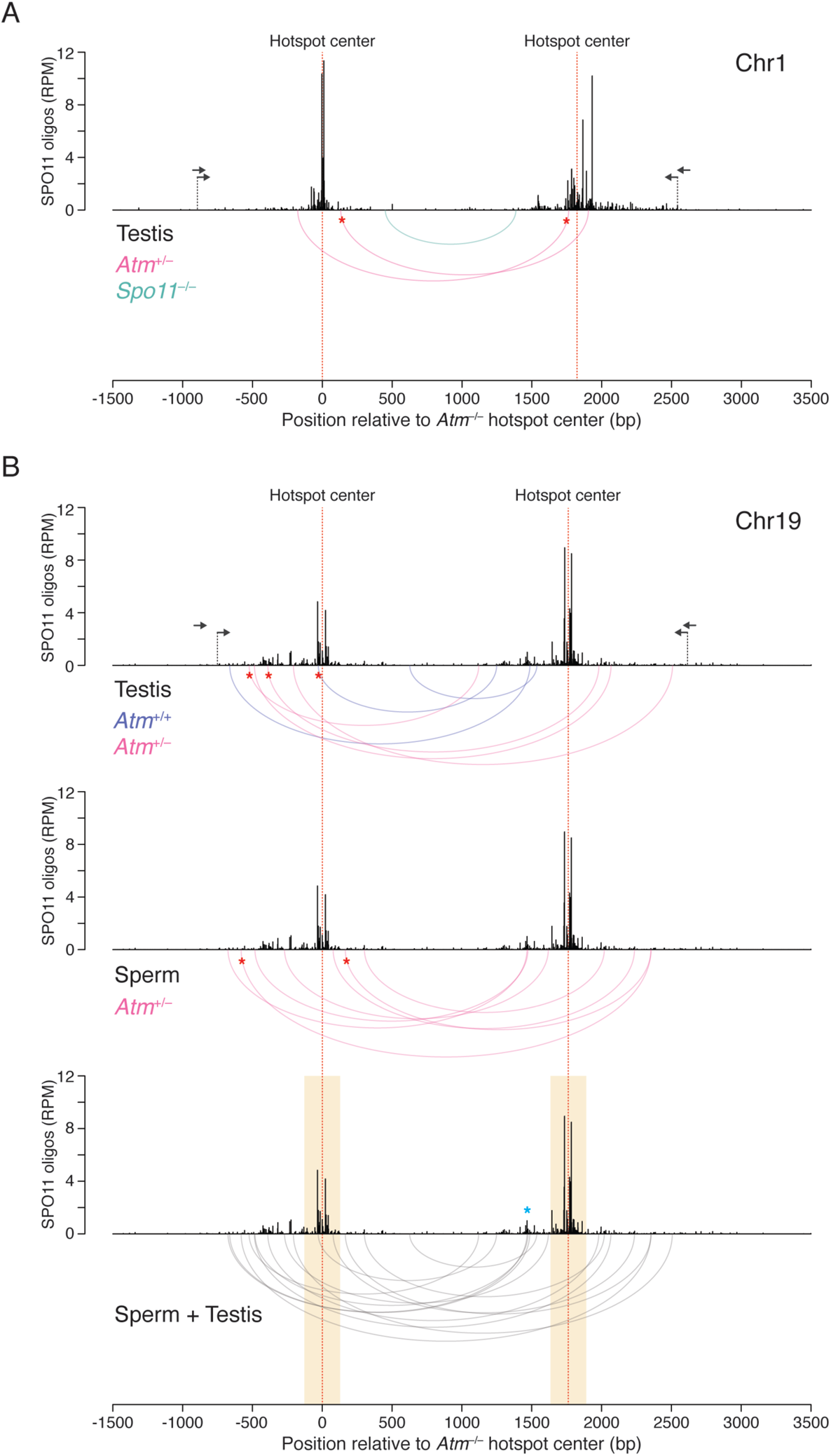
Deletions detected in the presence of ATM in testes and sperm. Deletions detected in *Atm*^+/+^ (blue arcs) and *Atm*^+/−^ (pink arcs) often had one breakpoint that was also found for a deletion in *Atm*^−/−^ (red asterisks). (A) For the Chr1 hotspot pair, two deletions were detected in *Atm*^+/−^ testis. The single deletion obtained from *Spo11*^−/−^ is also shown (green arc). (B) For the Chr19 hotspot pair, deletions from *Atm*^+/+^ and *Atm*^+/−^ testis and *Atm*^+/−^ sperm are plotted separately and together. Note that 12 of 14 deletions mapped outside of the 251-bp zone around the hotspot centers (yellow blocks), which is in contrast to deletions observed in *Atm*^−/−^ (**Fig. S3**). The blue asterisk indicates a breakpoint cluster overlapping a SPO11-oligo cluster.

**Figure S6.**
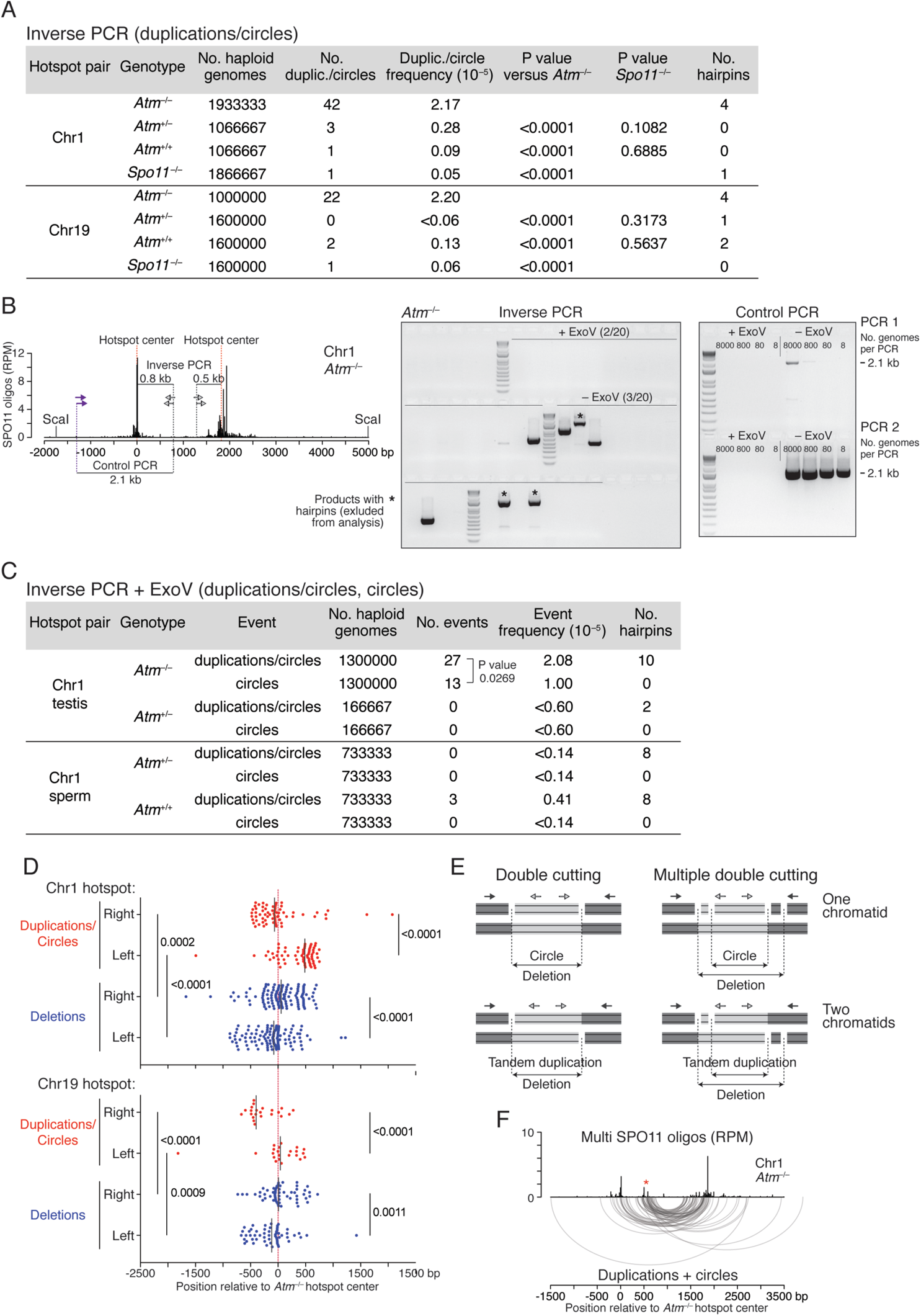
Extended analysis of tandem duplications and circular DNA. **(A)** Frequencies of duplications/circles from the inverse PCR experiments. Sequencing was used to verify duplication/circle products. The few products that could not be fully sequenced were excluded from the frequency calculation. Partial sequences indicated the presence of palindromes of ≥7–30-bp that would impede sequencing because of hairpin formation. These products were found in all genotypes, indicating likely PCR errors or SPO11-independent events. One mouse of each genotype was analyzed for both hotspot pairs. P values are from the Chi-square test. **(B)** ExoV experiment to detect circular DNA at the Chr1 hotspot pair, related to Fig. 3C**, D**. Left, SPO11-oligo map in *Atm*^−/−^ and the location of nested primer pairs (arrows) used in inverse PCR (light gray) and in control PCR (purple and light gray). Restriction sites for ScaI, which was used to digest genomic DNA prior to ExoV treatment, flank the hotspot pair. Middle, representative gel of inverse PCR products obtained from reactions either digested with ExoV to detect circles or not digested with ExoV to detect duplications/circles. Note that ExoV treatment appears to remove products that form hairpins. Right, corresponding gel of control PCR to assess ExoV efficiency, showing PCR products from the primary (PCR 1) and secondary (PCR 2) PCRs. The estimated number of pre-digestion haploid genomes per PCR is shown. No products were amplified in PCRs from reactions treated with ExoV. **(C)** Frequencies of duplications/circles and circles from the ExoV experiments at the Chr1 hotspot pair. For *Atm*^−/−^, testis DNA from two mice was analyzed. No products were observed in the small-scale experiment using *Atm^+/−^* testis DNA. In the larger experiment with *Atm^+/−^* and *Atm^+/+^* sperm from one mouse per genotype, three duplication/circle products were detected, but no circles. This would be consistent with the idea that circle DNA is not transmitted into sperm; however, their low frequency and the more frequent appearance of PCR products with hairpins in –ExoV experiments obscures our evaluation of these genotypes. P value is from the Chi-square test. **(D)** Distribution of breakpoint distances from the hotspot centers for deletions and duplications/circles, related to Fig. 3F. Scatter plots of breakpoint positions at the left and right hotspots in the Chr1 and Chr19 hotspot pairs. P values are from an unpaired Mann-Whitney test. **(E)** Multiple double cuts should shift breakpoint junctions for deletions versus tandem duplications and circles. Left, double cutting at two DSB hotspots (one DSB per hotspot) is expected to lead to similarly sized deletions, tandem duplications, and circles. Right, multiple double cuts involving additional DSBs at each hotspot will lead to shorter tandem duplications and circles, but larger deletions. **(F)** Imputed multi-mapping SPO11 oligos at the Chr1 hotspot pair (Lange et al., 2016). For the left hotspot, duplication/circle breakpoints frequently map at and close to a SPO11-oligo cluster at ~ 500 bp from the hotspot center (red asterisk), which is not observed in the unique SPO11-oligo map (Fig. 3F).

**Figure S7.**
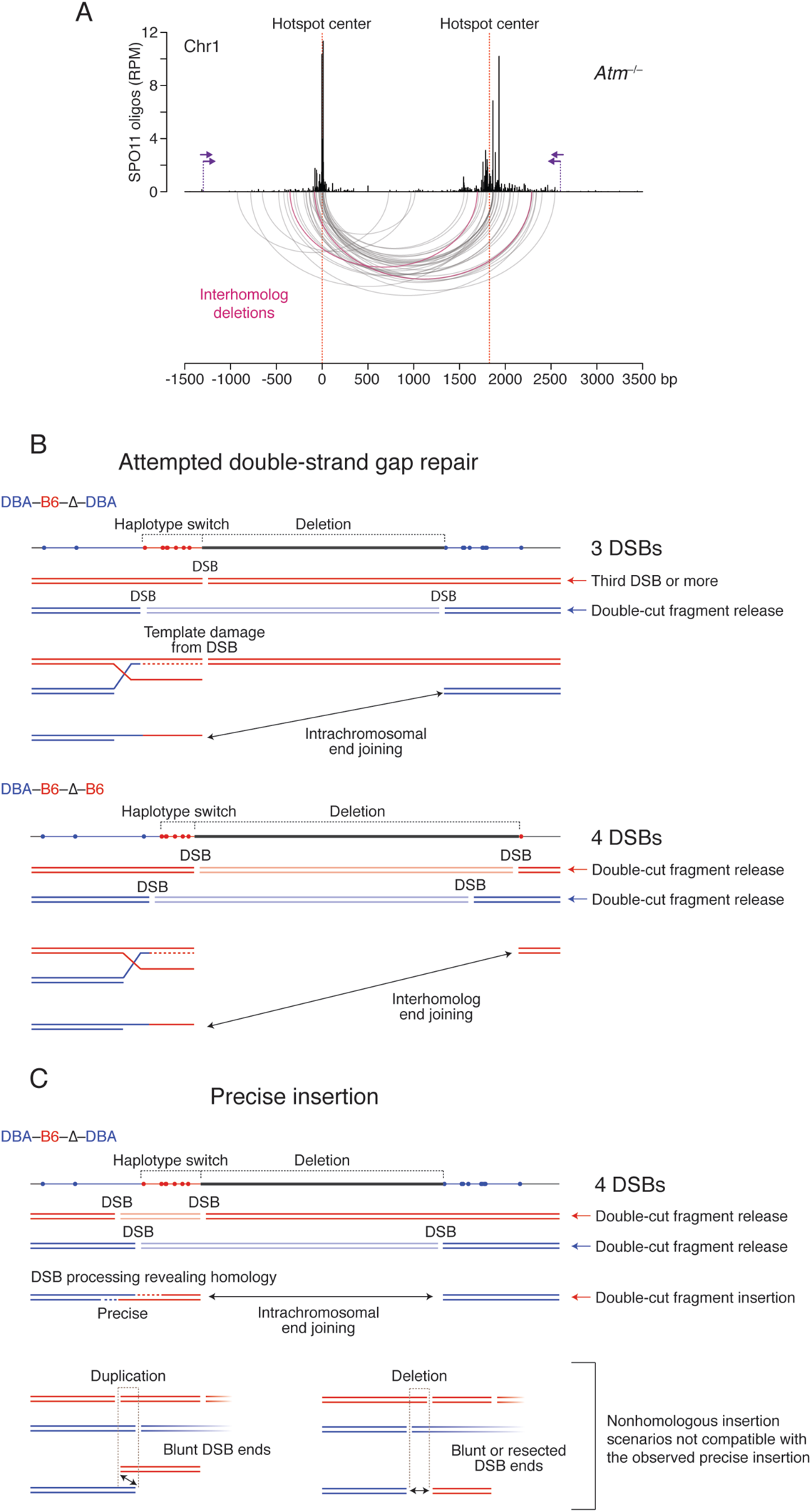
Deletions in *Atm*^−/−^ B6 × DBA F1 hybrid mice with models for haplotype switching. **(A)** Arc diagram of deletions from the Chr1 hotspot pair (37 deletions total). Two interhomolog deletions are indicated (pink arcs). **(B, C)**. Models for haplotype switching in deletions. **B**, Attempted double-strand gap repair. Examples of one intrachromosomal (top) and one interhomolog (bottom) deletion with haplotype switches. In both cases, two DSBs produce a double-stranded gap between the two hotspots, releasing a double-cut fragment (lighter blue or red double line). One DSB end at the gap invades the homolog and is extended by DNA synthesis, but a DSB in the repair template leads to aborted synthesis, such that this end is ejected and joins the other end of the gap (intrachromosomal end joining) or the end of a DSB formed at the other hotspot in the homolog (interhomolog end joining). In the case of interhomolog deletions, it is also possible that only one DSB is formed on the chromosome initiating strand invasion. Thus, in both examples at least three DSBs occur, indicating extensive damage to multiple chromatids at adjacent hotspots when ATM is absent. The substantial template damage in this scenario may favor nonhomologous end-joining mechanisms. **C**, Precise insertion due to homologous annealing. An example of an intrachromosomal deletion with a haplotype switch. At a minimum four DSBs are formed, involving double cutting within one hotspot on one chromosome and at both hotspots on the other chromosome. A released double-cut fragment from one hotspot is precisely inserted at the double-stranded gap in the homolog through annealing of homologous sequences exposed by resection. Importantly this scenario depends on the relative position of the two DSBs in chromosomes. If the relative position were switched (bottom right), resection would not expose homologous sequences. Nonhomologous insertion is not compatible because it would produce a small deletion (bottom right) or a small duplication (bottom left). As we do not observe these types of repair products in our deletions, we consider the attempted double-strand gap repair scenario more likely.

**Figure S8.**
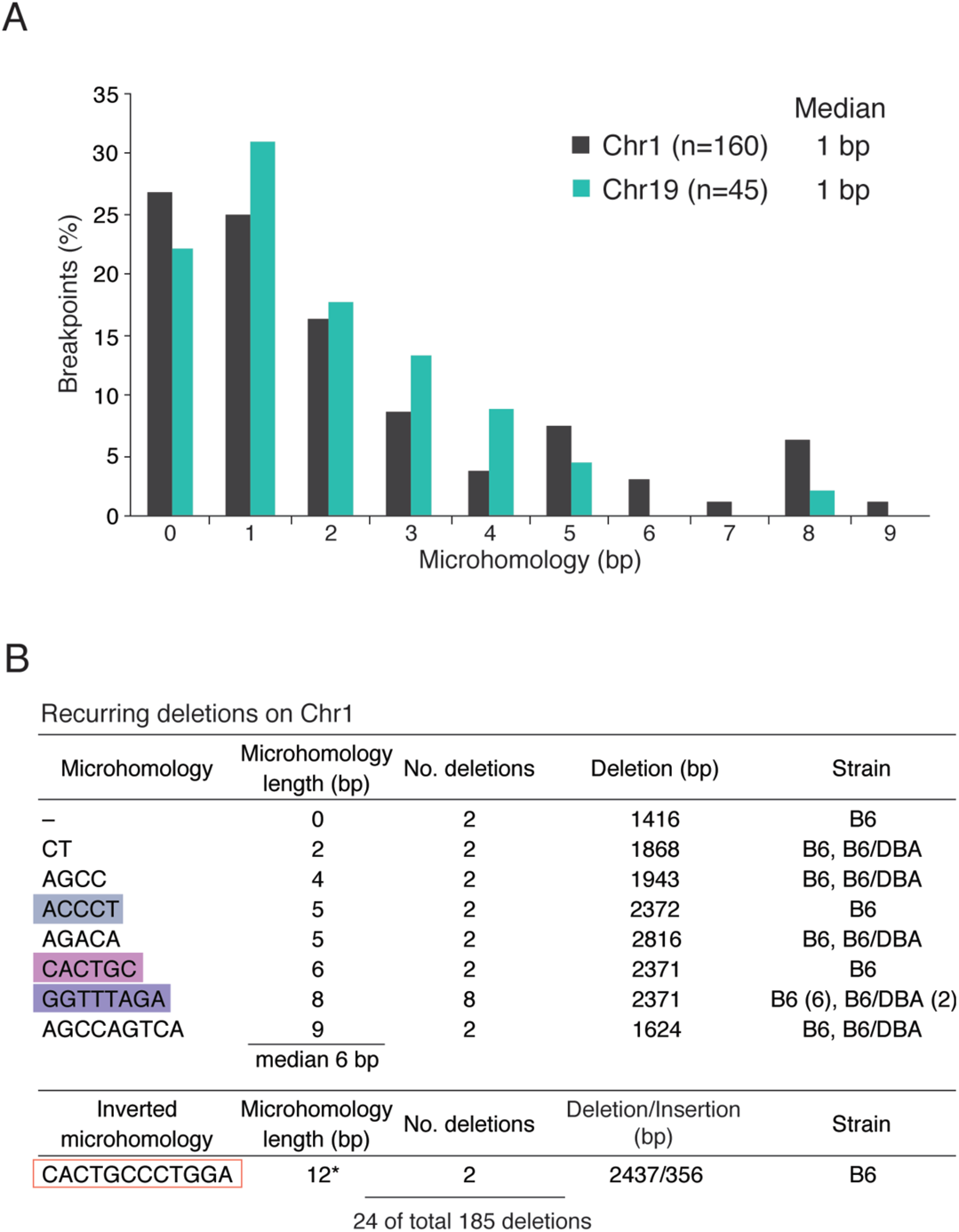
Microhomology at deletion breakpoints in *Atm*^−/−^. **(A)** Distribution of microhomology lengths for the Chr1 and Chr19 hotspot pairs deletion breakpoints. Only deletions without insertions or other DNA-end modifications are included (**Table S2**). **(B)** Recurring deletions at the Chr1 hotspot pair in B6 inbred and B6 × DBA F1 hybrid mice. Recurring deletions often had longer microhomologies at breakpoints. Microhomologies within the larger imperfect repeat described in the text are marked in different colors as in Fig. 4C. A recurring deletion with inverted insertion (asterisk) with a 12-bp microhomology at one insert breakpoint is also shown (**Fig. 4C, S10B**).

**Figure S9.**
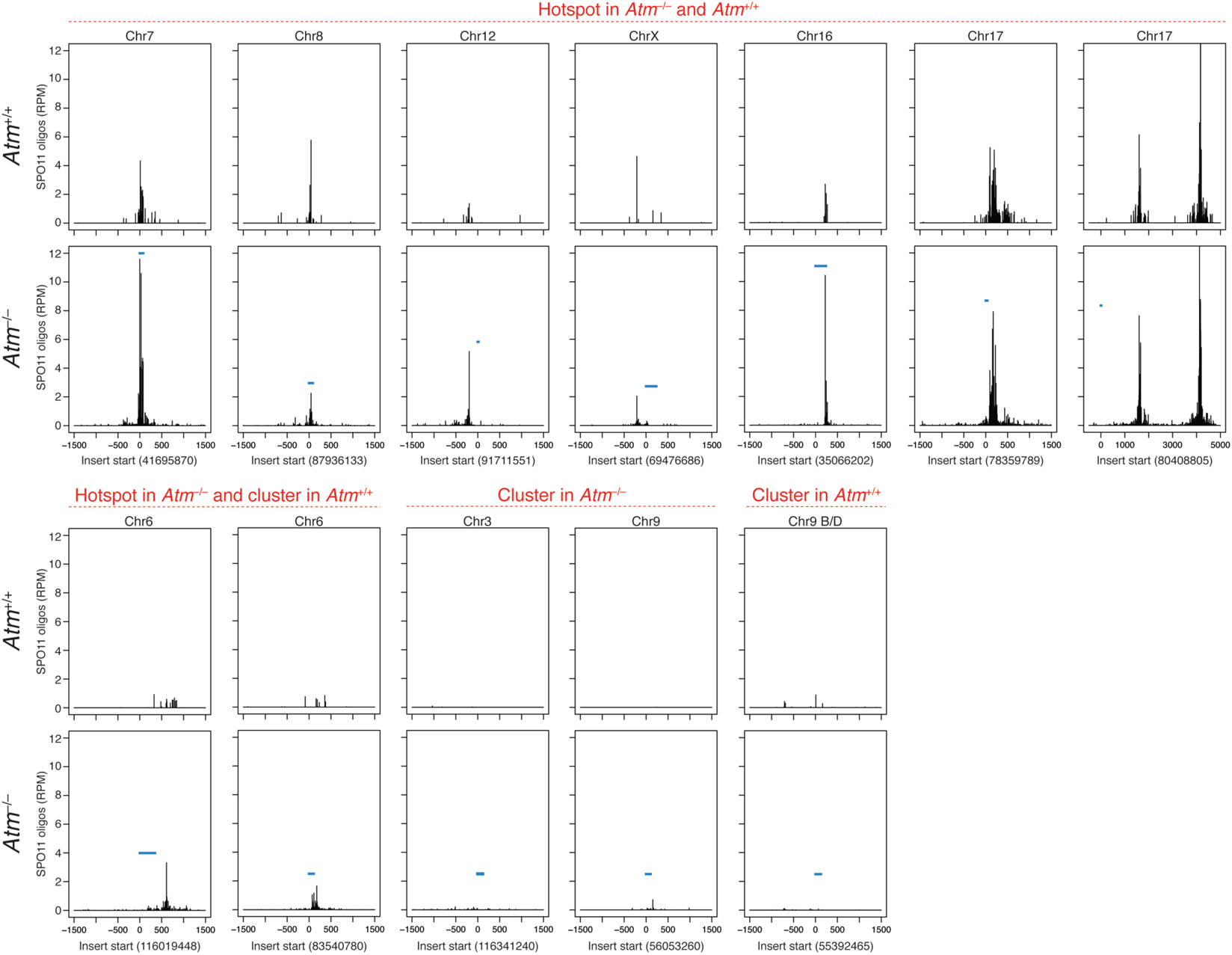
Derivation of inserts mapping to DSB hotspots on other chromosomes. SPO11 oligos at the genomic locations of inserts (blue bars) are shown for *Atm*^−/−^ and *Atm*^+/+^ and indicated as to whether they were called as hotspots in one or both genotypes or instead contain SPO11-oligo clusters. Inserts mapping to the non-PAR ChrY are shown in **Fig. S11**. One insert mapping to a microsatellite repeat is not shown. See also **Table S4**.

**Figure S10.**
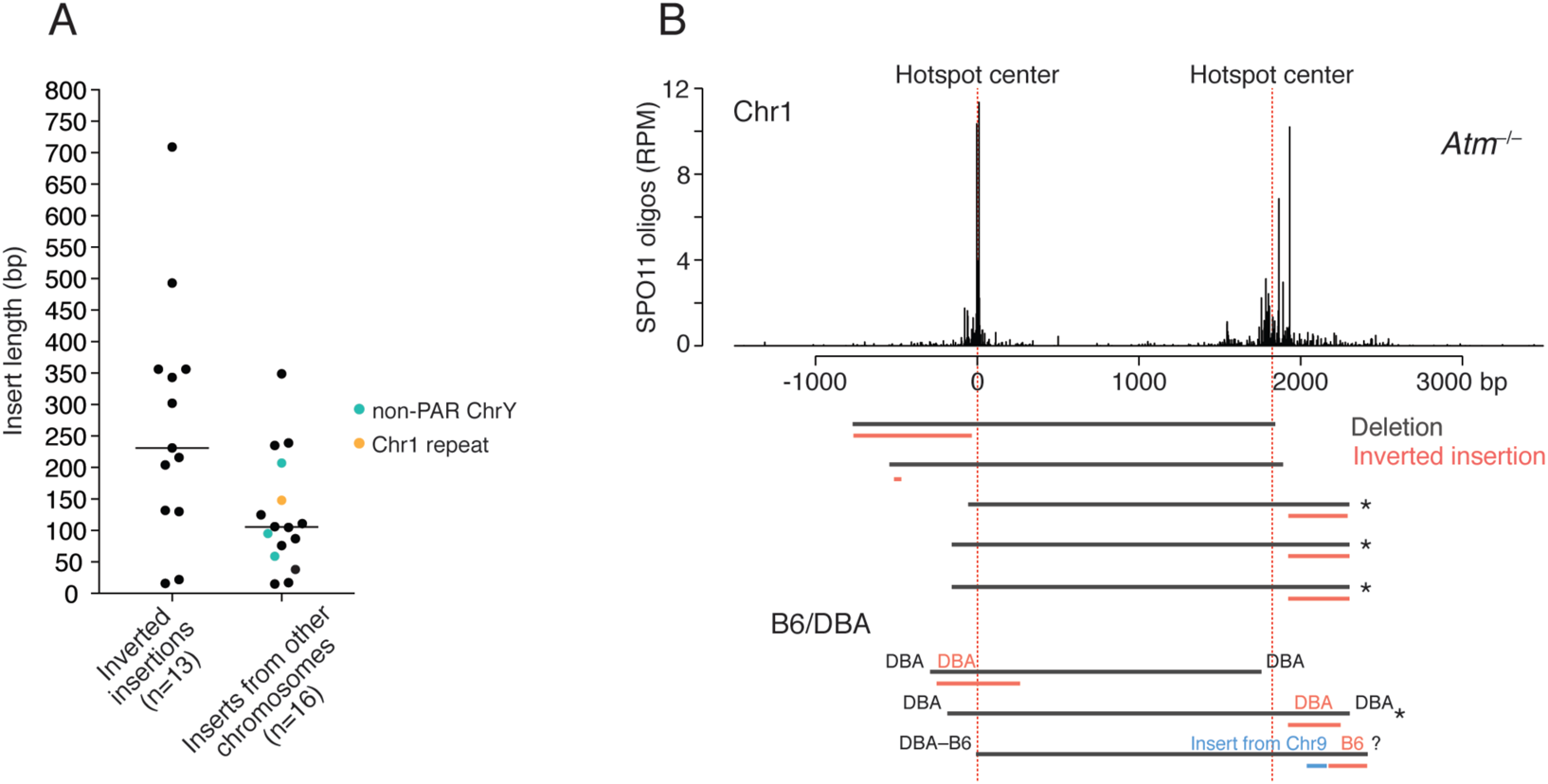
Characteristics of insertions in *Atm*^−/−^. **(A)** Inverted insertions from the same hotspot pair have a broader distribution of lengths than inserts derived from other chromosomes. Median insert lengths are indicated. See also **Tables S4, S5**. **(B)** Deletions with inverted insertions at the Chr1 hotspot pair in B6 and B6 × DBA mice. Asterisks denote deletions where one insert breakpoint had the same microhomology as the site with recurring deletions (Fig. 4C). Note that for the two deletions on the DBA chromosome, the inserts also derived from the DBA chromosome.

**Figure S11.**
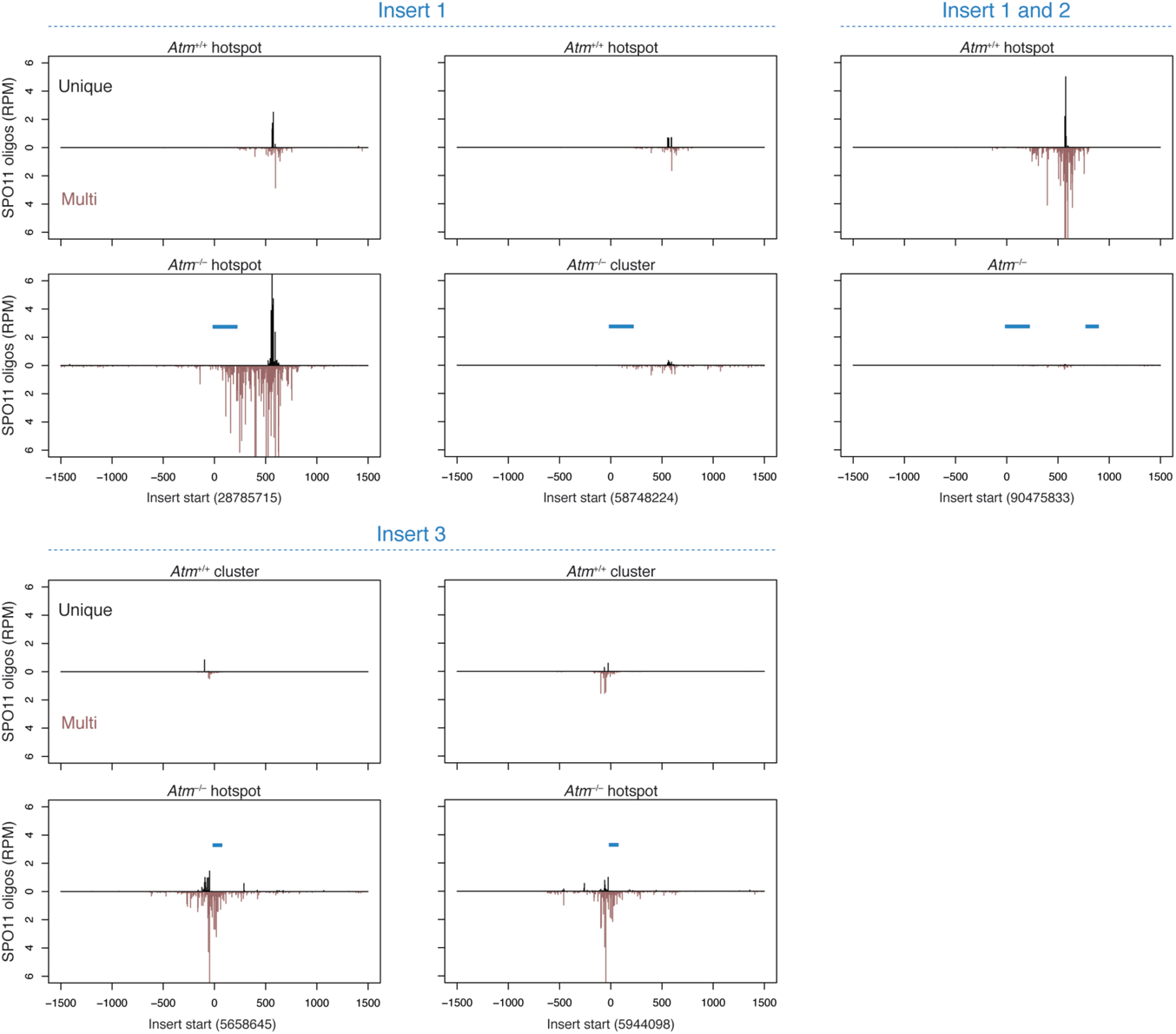
Inserts from the non-PAR ChrY map to multiple DSB hotspots. Three inserts identified in *Atm*^−/−^ deletions map to non-PAR ChrY repeats (**Table S4**), some of which have called hotspots. SPO11-oligo profiles are shown only for those repeats containing hotspots in *Atm*^−/−^, *Atm*^+/+^, or both, with the position of the inserts (blue bars) for each hotspot indicated. Inserts 1 and 2 are part of the same repeat family, and in one case they map to a single hotspot. Inserts 1 and 3 map to three and two hotspots, respectively. SPO11 oligos map uniquely to these hotspots (“Unique”) or also map to other copies of the repeats (“Multi”). It remains possible that other repeat locations to which SPO11 oligos do not map uniquely also contain hotspots. Given that all insert-matching repeats (67 total) contain an intact PRDM9 motif, it is possible that DSBs are formed in many such repeats. The locations of multi-mapping SPO11 oligos were imputed, meaning they were distributed among mapped positions proportionally to local densities of uniquely mapping SPO11 oligos (Lange et al., 2016).

**Figure S12.**
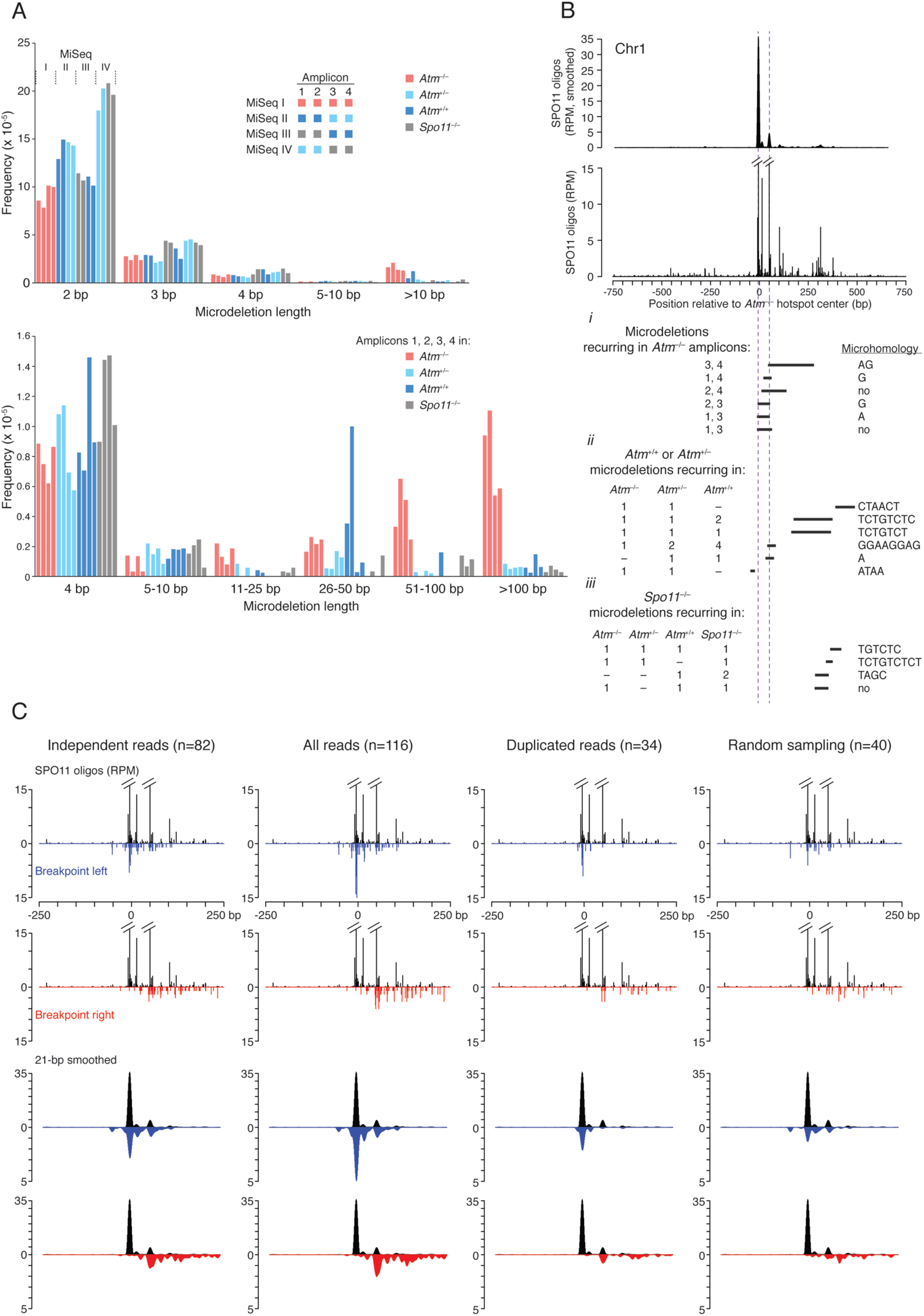
Evaluation of reads with microdeletions from deep sequencing of a single hotspot. **(A)** Frequency of reads that had microdeletions of the indicated lengths. Four overlapping amplicons across the hotspot (1–4) were deep sequenced using MiSeq. For each amplicon, 576 separate PCRs were carried out with a total input of 9.6 million haploid genome equivalents, using a limited number of amplification cycles (see Methods). The PCR products for each amplicon were then pooled for sequencing. Four MiSeq runs were performed (I–IV) with the indicated amplicons and genotypes. For clarity, sequencing results are grouped by run (Top) or genotype (Bottom). At top, note that reads with small deletions (2–4 bp) were frequently observed in all amplicons for each genotype, including *Spo11^−/−^*, and that their frequency varied between sequencing runs, indicating a significant batch effect. Short deletions (e.g., 2 bp) are known to arise from PCR and sequencing errors (Schirmer et al., 2015), and we saw that they became progressively less abundant in all genotypes up to 10 bp. Excluded from the 4-bp deletions is a product that was represented by >1000 reads in amplions 1–3. By contrast, reads with deletions >10 bp were more frequent in *Atm*^−/−^ than in the other genotypes. At bottom, note that reads with 11–25-bp microdeletions were more frequent in *Atm*^−/−^. Therefore, 11 bp was used as the lower cutoff to define a microdeletion for subsequent analyses. Our sequencing design (pooling of many low-cycle PCRs) is expected to minimize PCR jackpots and reduce instances where PCR duplicates from the same input DNA molecule are sequenced. However, two amplicons in *Atm*^+/+^ showed multiple reads for one microdeletion each (amplicon 1: 26-bp microdeletion, 14 reads; amplicon 2: 31-bp microdeletion, 22 reads; **Table S6**), which therefore may be PCR duplicates. Thus, in some subsequent analyses we applied a stringent definition of independent events by including only one read for a given deletion product from a given amplicon. **(B)** Recurring microdeletions within the hotspot. Top, smoothed (21-bp Hann filter) and unsmoothed SPO11-oligo maps at the Chr1 hotspot in *Atm*^−/−^. Bottom, recurring microdeletions. These identical microdeletions (black bars) were found in independent PCRs for different amplicons. i) In *Atm*^−/−^, amplicons that yielded the recurring microdeletions are indicated. For example, the microdeletion with the AG microhomology arose in amplicons 3 and 4. Note that one or both breakpoints for each of the recurring microdeletions occured at SPO11-oligo peaks (dashed vertical lines). ii) In *Atm*^+/−^ and *Atm*^+/+^, some microdeletions were identical to those found in *Atm*^−/−^, in some cases in multiple amplicons. For example, the microdeletion with the GGAAGGAG microhomology was recovered in all three genotypes, but in different amplicons (*Atm*^−/−^, 1; *Atm*^+/−^, 2; *Atm*^+/+^, 4). iii) Some of the microdeletions found in *Spo11*^−/−^ recurred in other genotypes. Note that these map to the same region of the hotspot away from the center. **(C)** Many of the duplicated reads in *Atm*^−/−^ amplicons likely represent independent microdeletion events rather than PCR duplicates. For each amplicon, we randomly sampled with replacement from a set of reads representing all of the unique microdeletions observed. The sampled group was equal in number to the duplicated reads actually obtained for that amplicon. Then, the sampled reads with microdeletions mapping to the region where amplicons 1, 2, and 3 overlap were combined, and their breakpoint distribution was compared with the distribution of the observed duplicate reads. Breakpoint profiles of independently obtained reads, as in Fig. 6C, and of all reads are also shown. In the smoothed breakpoint profiles of the observed duplicates, the left breakpoints were particularly enriched in a peak that matched the central peak of SPO11 oligos, and the right breakpoints were enriched at a position that matched the secondary peak of SPO11 oligos. These patterns were not recapitulated by random sampling: both the left and right breakpoints were more widely dispersed without such clear enrichment coincident with the SPO11-oligo peaks. Because the observed duplicates were more likely than the random sample to have breakpoints aligned with the strongly favored locations of DSBs, these findings are consistent with the conclusion that most of the duplicated reads represent independent microdeletion events.

**Figure S13.**
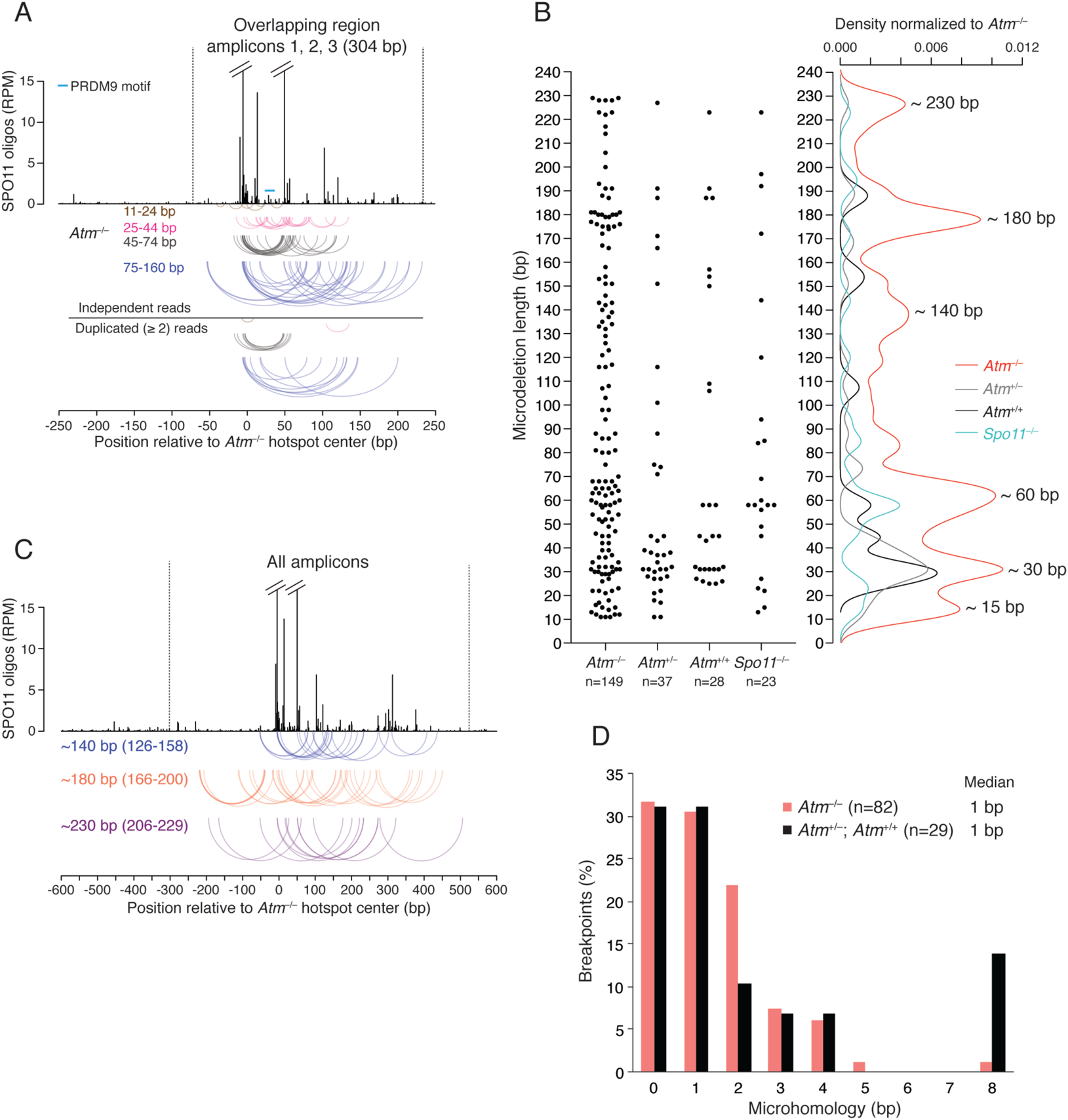
Extended analysis of microdeletions by genotype and microhomology. **(A)** Arc diagrams of microdeletions stratified by length from *Atm*^−/−^, as in Fig. 6C, with microdeletions in duplicated reads below. Microdeletions >160 bp are not shown, as their location may be more constrained by restricting analysis to the region where the amplicons overlap. **(B)** Microdeletion length distribution for all amplicons by genotype for independent reads. Scatter plots and corresponding density plots are shown (>10 bp); densities in *Atm*^+/−^, *Atm*^+/+^, and *Spo11*^−/−^ are normalized to *Atm*^−/−^ to reflect read numbers in each genotype. As in the analysis of microdeletions in the overlap region of amplicons 1, 2, and 3 (Fig. 6D), distinct size classes were observed in the presence and absence of ATM, but not in *Spo11*^−/−^. Larger microdeletion size classes (~140 bp, ~180 bp, ~230 bp) were well defined when all amplicons are considered. A smaller size class (~15 bp) was more prominent outside of the overlap region. **(C)** Arc diagrams of the ~140-bp, ~180-bp, and ~230-bp microdeletions observed for *Atm*^−/−^. **(D)** Microhomology lengths at microdeletion breakpoints in the presence and absence of ATM. Analyses were performed for independent microdeletions in the overlapping region between amplicons 1, 2, and 3.

**Table S1.**
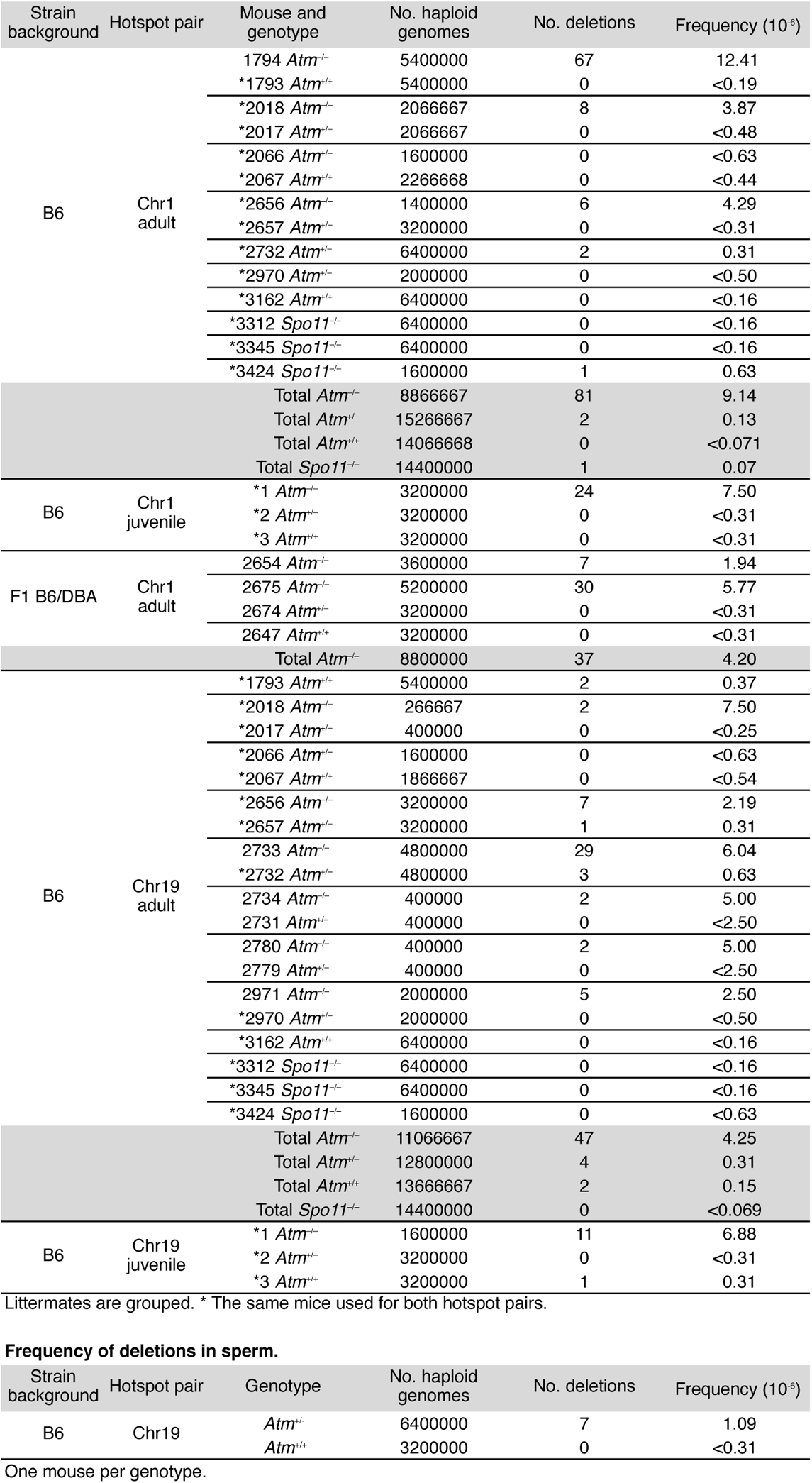
Frequency of deletions in testis from individual mice.

**Table S2.**
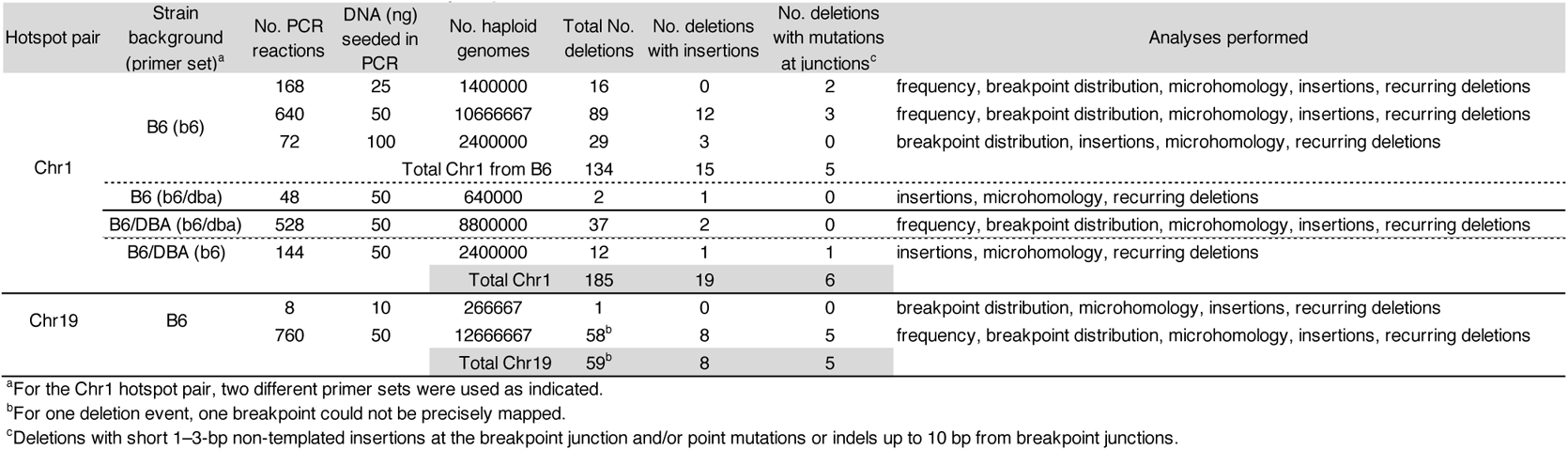
Deletions detected in Atm^−/−^ with the analyses performed.

**Table S3.** Comparison of expected double cutting frequency with observed deletion frequency in Atm^−/−^.

| Hotspot pair |  | Hotspot strength (RPM) | No. DSBs per chromatid at hotspot (DSB frequency per meiosis)<sup>a</sup> | Expected double cutting frequency per meiosis | Deletion frequency per meiosis<sup>b</sup> | Expected double cutting frequency divided by deletion frequency |
| --- | --- | --- | --- | --- | --- | --- |
| Chr1 | Hotspot left | 53 | 0.03975 | $27.7 \times 10^{-4}$ | $0.36 \times 10^{-4}$ | 77 |
|  | Hotspot right | 93 | 0.06975 |  |  |  |
| Chr19 | Hotspot left | 41 | 0.03075 | $15.9 \times 10^{-4}$ | $0.17 \times 10^{-4}$ | 94 |
|  | Hotspot right | 69 | 0.05175 |  |  |  |

<sup>a</sup>DSB frequency was estimated from SPO11-oligo counts (RPMs) by equating 3000 DSBs in *Atm*<sup>-/-</sup> to one million reads. Note: we considered that there are about 300 DSBs per meiotic cell in wild type and an ~10-fold increase in *Atm*<sup>-/-</sup>.
<sup>b</sup>Deletion frequency calculated per meiotic genome.

**Table S4.**
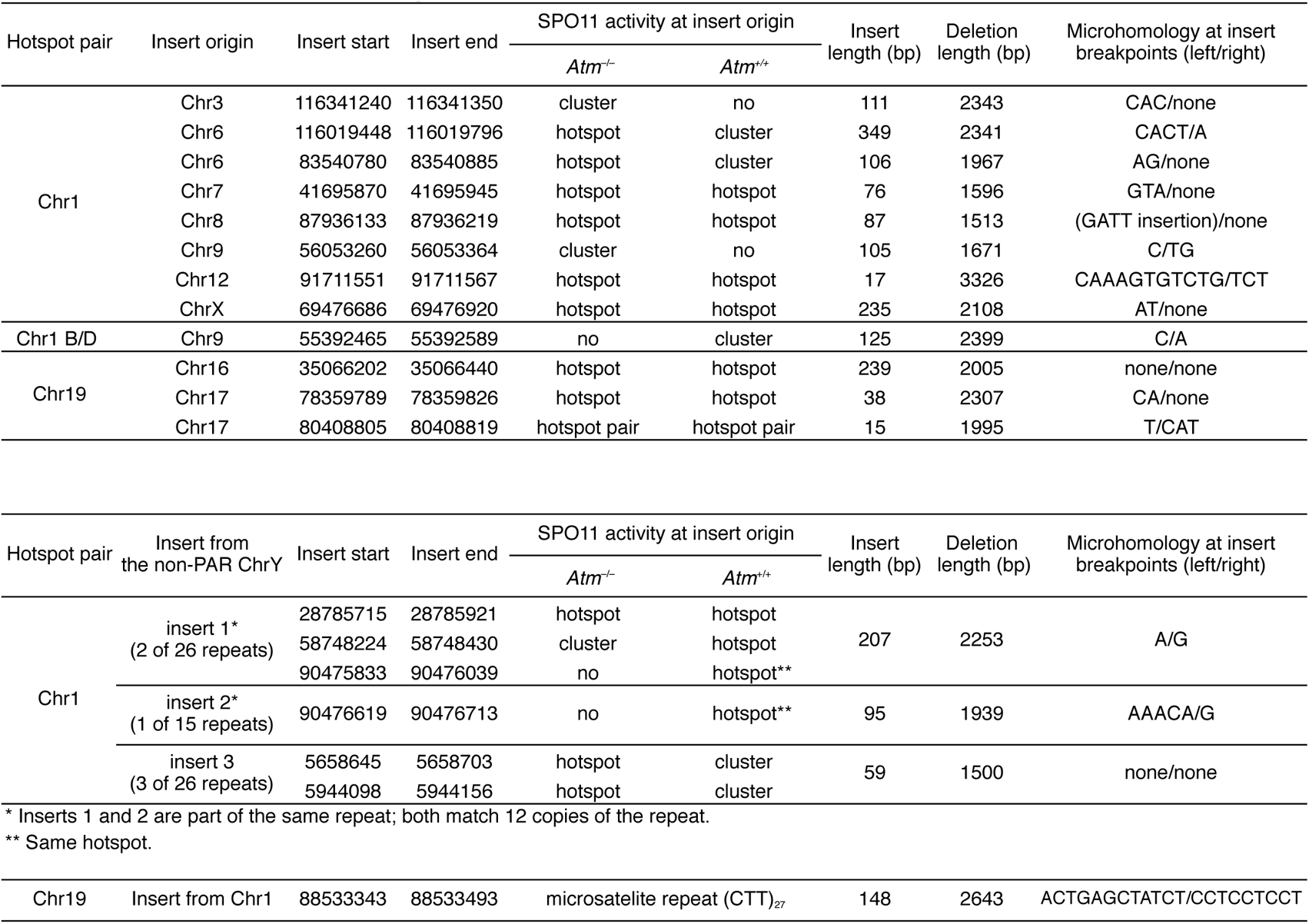
Insertions derived from other hotspots.

**Table S5.** Insertions derived from the hotspot pair.

| Hotspot pair | Hotspot left<br>or right | Insert<br>length (bp) | Deletion<br>length (bp) | Distance between<br>deletion and insertion<br>breakpoints (bp) | Microhomology<br>at insert breakpoints<br>(left/right) |
| --- | --- | --- | --- | --- | --- |
| Chr1 | left | 709 | 2590 | 4 | none/GAAGG |
|  | left | 22 | 2411 | 28 | none/AGA |
|  | right | 343 | 2334 | 11 | TCCAGGGCAGTGC/none |
|  | right | 356 | 2437 | 0 | TCCAGGGCAGTG/none |
|  | right | 356 | 2437 | 0 | TCCAGGGCAGTG/none |
| Chr1 B/D | left | 493 | 2025 | 40 | TTG/AAAG |
|  | right | 302 | 2465 | 56 | TCCAGGGCAGT/ATTC |
|  | right | 216 | 2399 | 3 | GT/T |
| Chr19 | left | 204 | 2591 | 0 | none/C |
|  | right | 231 |  | 3 | CTCTCTC/none |
|  | left | 130 | 1361 | 0 | none/ATAGCT |
|  | left | 132 | 1949 | 0 | none/TAGCT |
|  | right | 16 | 2063 | 44 | GCTCT/A |

**Table S6.**
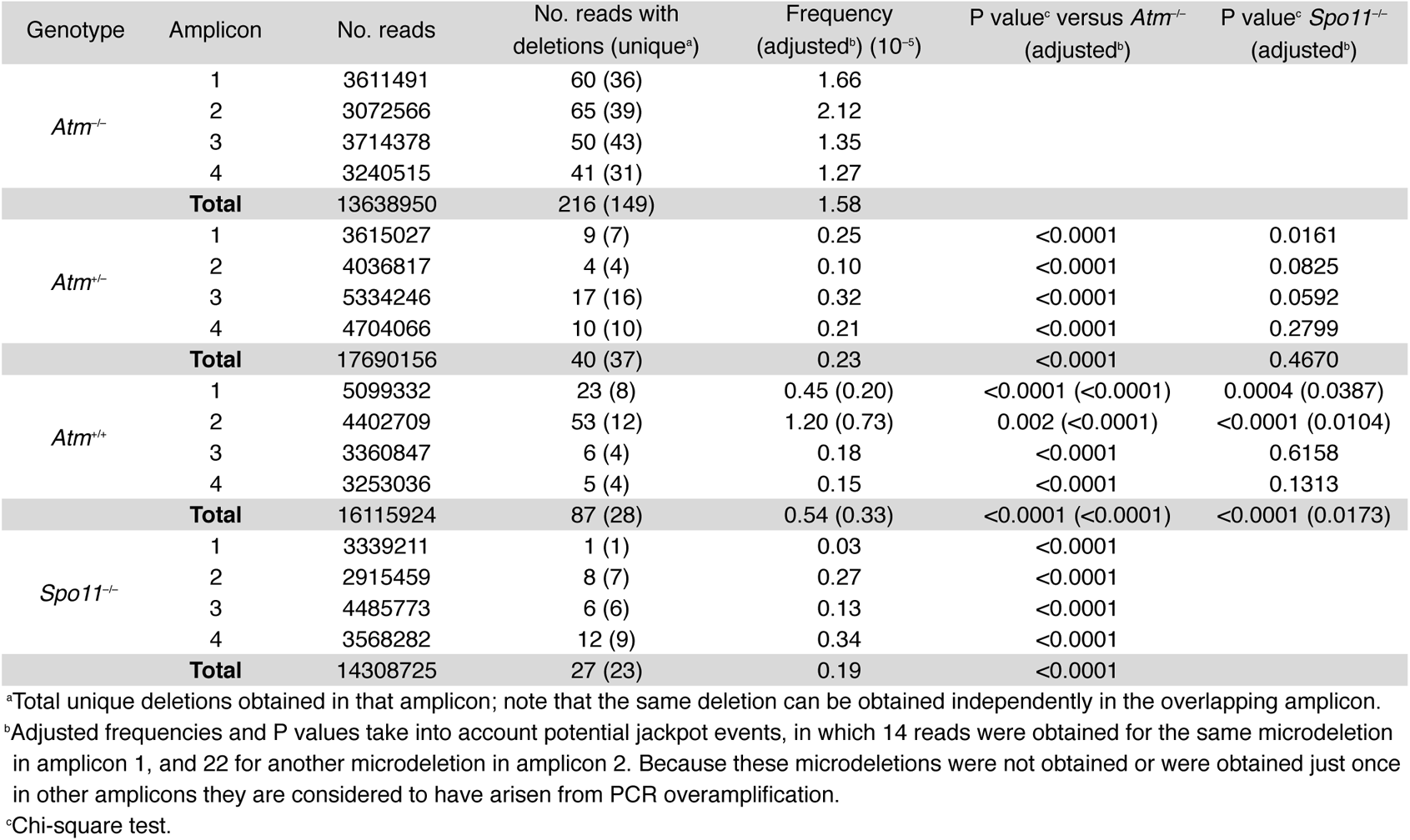
Frequency of microdeletions in individual amplicons.

